# Confinement and Catalysis Within *De Novo* Designed Peptide Barrels

**DOI:** 10.1101/2024.08.22.609140

**Authors:** Rokas Petrenas, Olivia A. Hawkins, Jacob F. Jones, D. Arne Scott, Jordan M. Fletcher, Ulrike Obst, Lucia Lombardi, Fabio Pirro, Graham J. Leggett, Thomas A.A. Oliver, Derek N. Woolfson

**Author notes:** Corresponding Authors Thomas A.A. Oliver, Derek N. Woolfson.

## Abstract

*De novo* protein design has advanced such that many peptide assemblies and protein structures can be generated predictably and quickly. The drive now is to bring functions to these structures, for example, small-molecule binding and catalysis. The formidable challenge of binding and orienting multiple small molecules to direct chemistry is particularly important for paving the way to new functionalities. To address this, here we describe the design, characterization, and application of small-molecule:peptide ternary complexes in aqueous solution. This uses α-helical barrel (αHB) peptide assemblies, which comprise 5 or more α-helices arranged around central channels. These channels are solvent accessible, and their internal dimensions and chemistries can be altered predictably. Thus, αHBs are analogous to ‘molecular flasks’ made in supramolecular, polymer, and materials chemistry. Using Förster resonance energy transfer as a readout, we demonstrate that specific αHBs can accept two different organic dyes, 1,6-diphenyl-1,3,5-hexatriene and Nile Red in close proximity. In addition, two anthracene molecules can be accommodated within an αHB to promote photocatalytic anthracene-dimer formation. However, not all ternary complexes are productive, either in energy transfer or photocatalysis, illustrating the control that can be exerted by judicious choice and design of the αHB.

## 1. INTRODUCTION

Natural enzymes catalyze chemical reactions with high substrate specificity, product stereoselectivity, and substantial accelerations of reaction rates. Designable and tunable enzyme-like catalysts would have wide-ranging applications in basic science, chemical synthesis, and the biotech and pharmaceutical industries. Despite considerable advances in the past four decades^1-6^, truly *de novo* peptide and protein design for small-molecule binding and efficient catalysis remains a significant challenge.^7-9^

Early *de novo* structural design used straightforward patterning of hydrophobic and polar residues to deliver peptide assemblies that mimicked simple protein architectures.^1, 3^ Some of these have been embellished with metal-binding sites leading to catalysis.^3, 10, 11^ These minimal approaches gave way to rational design, in which sequence design was augmented by sequence-to-structure/function relationships garnered from natural proteins to produce a wider variety of robust peptide and protein designs.^1, 12^ Such designs have also been functionalized to yield hydrolases and various metalloenzymes amongst other activities.^11, 13-17^ In parallel with these developments, computational protein design emerged to deliver methods for backbone constructions, for fitting *de novo* sequences onto these scaffolds, and for assessing the quality of the models ahead of experiments.^11, 18^ Early computational protein design also introduced modelling of protein-ligand interactions.^19^ This led to natural protein scaffolds being repurposed for binding and catalysis.^20-22^ Currently, the field is undergoing another step-change with the application of deep-learning methods to generate *de novo* protein sequences, structures, and functions.^5, 23-25^ This has allowed the design of complex structures with tailored binding functions, in some cases with nanomolar affinity and sub-Å accuracy.^7, 8, 26^ However, overall success rates of completely *de novo* binders and enzymes remain low and usually require screening of large libraries and/or further optimization *via* directed evolution.^6-9^

Despite this trajectory and advances in rational and computational design, there are very few successful examples of bringing two or more ligands together into a fully *de novo* designed protein binding site.^27, 28^ However, with many cofactor-like catalysts available^29^, reactions in aqueous solutions could be accelerated by confining and orienting multiple small molecules within minimal binding sites to direct chemistry between them.

Although catalysis by confinement is observed in natural enzymes^30^, the concept of “molecular flasks” have been exploited broadly in supramolecular and polymer chemistry, where reactions in aqueous solution are accelerated by encapsulating guest molecules in micelles^31^, organometallic cages^32, 33^ or ordering of molecules by crystallization.^34, 35^ Perhaps the most biomimetic are the organometallic cages, as the shape and size of a hydrophobic pocket controls ligand binding and regio- and stereo-selectivity of a reaction, including new reactions not possible in free solution.^32, 33^ Examples of reactions catalyzed by these cages include regioselective 1,3-dipolar cycloadditions, and a novel Diels-Alder reactivity between inert arenes and N-substituted maleimides.^32^

In the last decade, a range of oligomeric α-helical barrels (αHBs) have been designed based on self-assembling peptides.^36-38^ Similarly to molecular flasks^39^, these αHB assemblies have solvent-accessible lumens that can bind small molecules, including reporter fluorescent dyes such as 1,6-diphenyl-1,3,5-hexatriene (DPH) and prodan.^37, 40^ The αHBs present interesting *de novo* scaffolds because of their stability, controllable oligomeric states and lumen sizes, robustness to mutation, and the potential to functionalize the internal channels for small-molecule binding and catalysis by incorporating both proteinogenic and non-canonical amino acids.^13, 41^

Inspired by the simplicity of molecular flasks, here we explore if αHBs can controllably bind two different small molecules and orient them for catalysis. By combining ultrafast spectroscopic methods to monitor Förster resonance energy transfer (FRET) and molecular dynamics simulations, we demonstrate that specific αHBs can accommodate two different organic dyes, DPH and Nile Red in close proximity. In addition, two anthracene molecules can be bound within the αHBs to promote or inhibit photocatalytic anthracene-dimer formation.

## 2. RESULTS

### 2.1 αHBS CAN ENCAPSULATE PAIRS OF SMALL MOLECULES

The lengths and widths of the channel in hexameric and heptameric αHBs (∼46 x 7 Å and ∼46 x 8 Å, respectively) are larger than ligands such as DPH (∼14 x 4 Å).^37^ Therefore, we hypothesized that it should be possible to bind multiple copies of such ligands within a single channel simultaneously, and potentially in conformations leading to productive interactions. We chose to probe the ternary complex formation through FRET. DPH was chosen as a potential FRET donor as it binds to multiple αHBs with µM affinities, characterized by a large increase in DPH fluorescence intensity.^41^ However, none of the dye molecules studied to date that bind αHBs had appreciable overlapping absorption with DPH emission and, thus, were not suitable as FRET acceptors.^37, 41^ Therefore, we used docking to identify three potential DPH-FRET acceptors with favorable characteristics for αHB binding; namely, Methyl orange, Coumarin-7, and Nile Red (Figure S3).^42^ Experimentally, both Coumarin-7 and Nile Red showed FRET when incubated with a heptameric αHB (CC-Type2-[I_a_V_d_], PDB code 6g66; Figure 1). Nile Red was selected as a FRET acceptor for further study as its emission was most intense and its fluorescence maxima significantly red-shifted from the peak of DPH’s emission (Figure 1).

**Figure 1.**
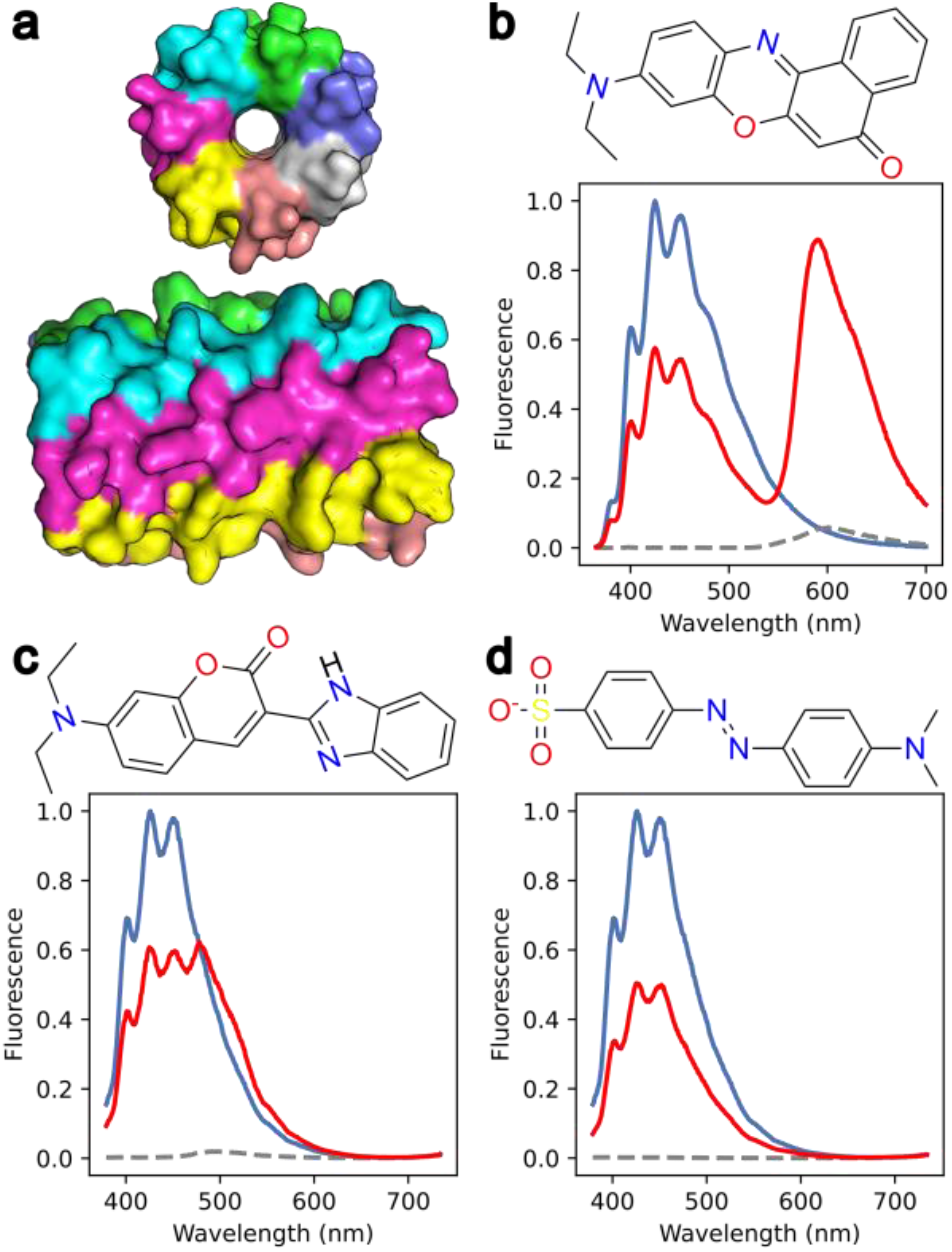
Nile Red and Coumarin-7 are FRET acceptors with DPH donor in the presence of an αHB. (a) Surface representation of the heptamer αHB crystal structure (PDB code 6g66). Colored by chain. (b – d) Steady-state emission spectra after DPH excitation within the heptamer with (b) Nile Red, (c) Coumarin-7 and (d) Methyl orange. Conditions: λ_ex_ = 352 nm, 3 μM potential acceptor 3 μM DPH, 5 μM peptide assembly, HEPES, 10% v/v MeCN, pH 7. Key: DPH emission spectrum, blue line; acceptor emission spectrum, broken black line; mixed DPH + acceptor emission spectrum, red line.

With this potential FRET pair in hand, we screened a wider set of αHBs for their Nile Red binding affinity and FRET efficiency. We restricted the screen to 15 open αHBs with hexameric to octameric oligomeric states and lumenal hydrophobic and/or aromatic residues (Table S1).^41^ As a negative control, we included a 3-helix bundle without a channel (PDB code 4nzl).^37^ This screen confirmed Nile Red binding to all 15 αHBs through steady-state emission spectra. Like DPH, the Nile Red fluorescence intensity was significantly increased when incubated with the αHBs as compared to the control (see Figures S4 and S5).

The screen identified 10 peptides that showed observable FRET between DPH and Nile Red (Figure 2a, Figure S6), and we selected the two αHBs with different oligomeric states that generated strong apparent steady-state FRET signals for further characterization: heptameric CC-Type2-[I_a_V_d_] and octameric CC-Type2-[I_a_I_d_]-I17V-I21F (PDB codes 6g66 and 9f5a). Both barrels bound DPH and Nile Red with low µM affinities. 2D excitation–emission data showed that FRET occurred from all vibronic states of the lowest energy DPH absorption band (Figure 2b,c), as evident from the strong ‘cross-peaks’ in the upper left quadrants of the correlation maps.

**Figure 2.**
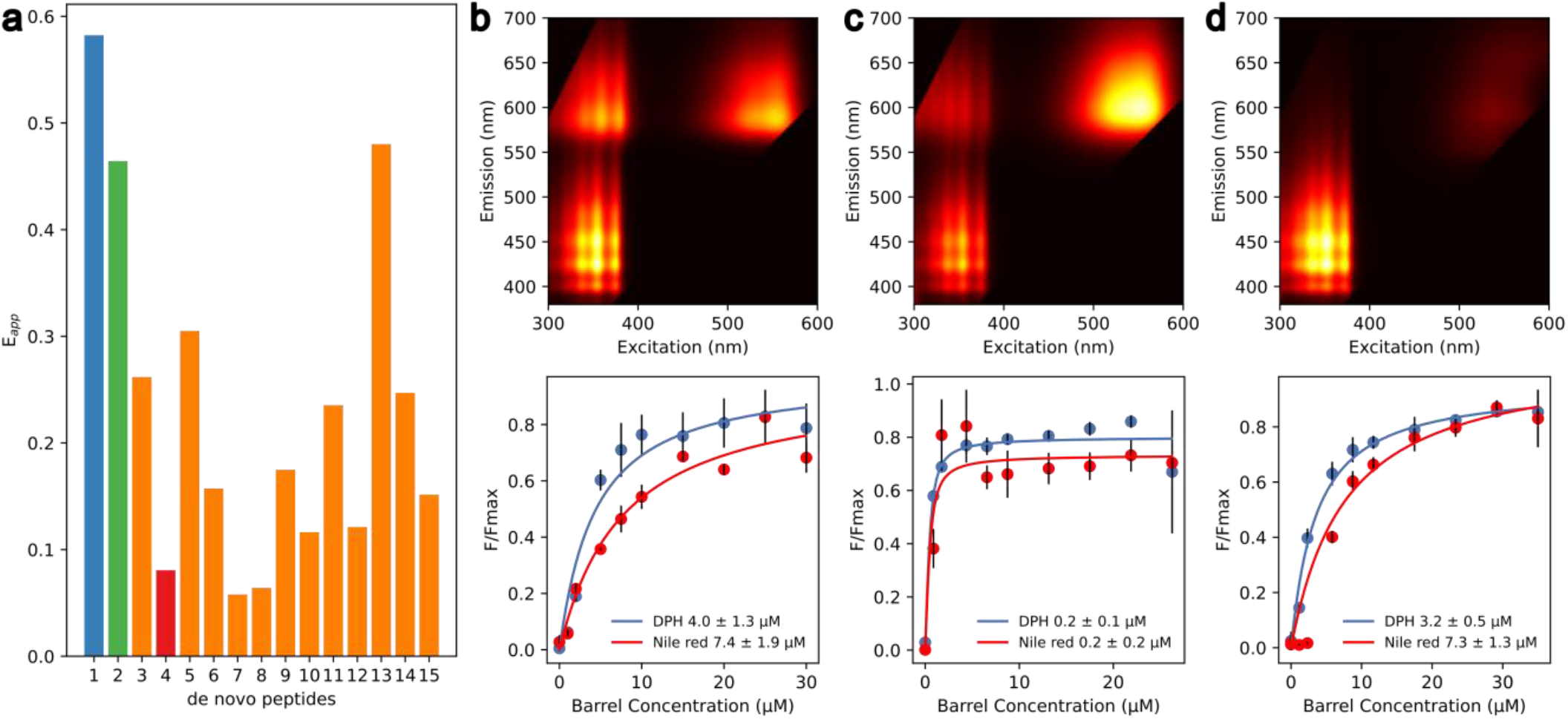
Screening for αHBs that co-encapsulate DPH and Nile Red. (a) Apparent steady-state FRET efficiency for the 15 screened peptides; the heptamer is in blue (#1), the octamer is green (#2) and the hexamer is red (#4), where 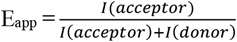. 2D excitation-emission spectra and the saturation binding curves for the top two hits: (b) the heptamer, (c) the octamer, and (d) a negative hit, the hexamer. F/F_max_: normalized fluorescence at 450 nm (DPH, blue) or 593 nm (Nile Red, red). Fluorescence conditions: 3 μM Nile Red, 3 μM DPH, 5 μM peptide assembly. Saturation binding curve conditions: 0.5 μM DPH or Nile Red, 0 – 30 μM peptide assembly, data are the mean of three independent repeats, error bars represent the standard deviation from the mean. All data collected in HEPES, 10% v/v MeCN, pH 7. See Table S1 for peptide biophysical data and sequences. See Figure S6 for the steady-state FRET spectra.

As a control for subsequent experiments described below, we chose a hexameric αHB (CC-Type2-[S_g_L_a_I_d_]; PDB code 4pn9) that bound both DPH and Nile Red individually with µM affinities but did not exhibit FRET (Figure 2d). In this case, DPH could displace Nile Red from the αHB, as evident from the decrease in Nile Red fluorescence intensity (Figure S8). This barrel exemplifies how cavity shape and size alone can lead to high specificity between two ligands with similar physical and chemical properties.

For the sake of simplicity, henceforth we refer to the three αHB peptides CC-Type2-[S_g_L_a_I_d_], CC-Type2-[I_a_V_d_] and CC-Type2-[I_a_I_d_]-I17V-I21F, as the hexamer, heptamer and octamer, respectively.

### 2.2 αHBS INCREASE ENERGY TRANSFER RATE BETWEEN DPH AND NILE RED

To characterize the interaction between DPH and Nile Red in αHBs further, we measured the timescales of energy transfer between the donor (DPH) and acceptor (Nile Red) dyes as these are highly sensitive to the intermolecular separation distance. This used time-correlated single-photon counting (TCSPC) and transient absorption (TA) spectroscopy performed on ternary complexes with the heptamer and the octamer αHBs.

First, we measured the fluorescence lifetimes for each dye individually in the barrels. Direct excitation of DPH at 352 nm yielded average fluorescence lifetimes of 15.9 ± 0.2 ns with the heptamer, and 15.8 ± 0.2 ns with the octamer (Table S4, Figure S12). For Nile Red, the corresponding control measurements used 535 nm light to selectively excite the dye. This yielded fluorescence lifetimes of 4.10 ± 0.2 ns (the heptamer) and 4.2 ± 0.2 ns (the octamer) (Table S5, Figure S14). In the presence of DPH, these increased to 4.9 ± 0.2 ns and 5.1 ± 0.2 ns, respectively (Figure S14, Table S6). This increase in Nile Red’s fluorescence lifetime (and small changes in the steady-state spectra, Figure S9) suggests that DPH alters the binding environment of Nile Red in the αHBs. Overall, when bound to the αHBs, the fluorescence lifetimes of DPH and Nile Red are at least as long as those previously reported in non-polar solvents, indicative of binding within enclosed hydrophobic environment.^43-45^

Next, we measured the timescale for energy transfer between DPH and Nile Red after direct and *selective* photoexcitation of DPH at 352 nm. Analysis of these data yielded FRET time constants of 0.98 ± 0.2 ns for the heptamer and 1.3 ± 0.2 ns for the octamer (Figures 3a and S17). Using the Förster equation (eq. S6), the inter-nuclear distance between DPH and Nile Red was estimated at 2.9 ± 0.1 nm in both αHBs (Figure S19). This is well within the length of the hydrophobic channels at ∼4.6 nm and demonstrates successful encapsulation of DPH and Nile Red within the same barrel. As anticipated from steady-state fluorescence experiments (Figure 2d), no evidence for FRET was observed in the hexamer from TCSPC measurements (Figures S12 – S15, Tables S4 – S6).

**Figure 3.**
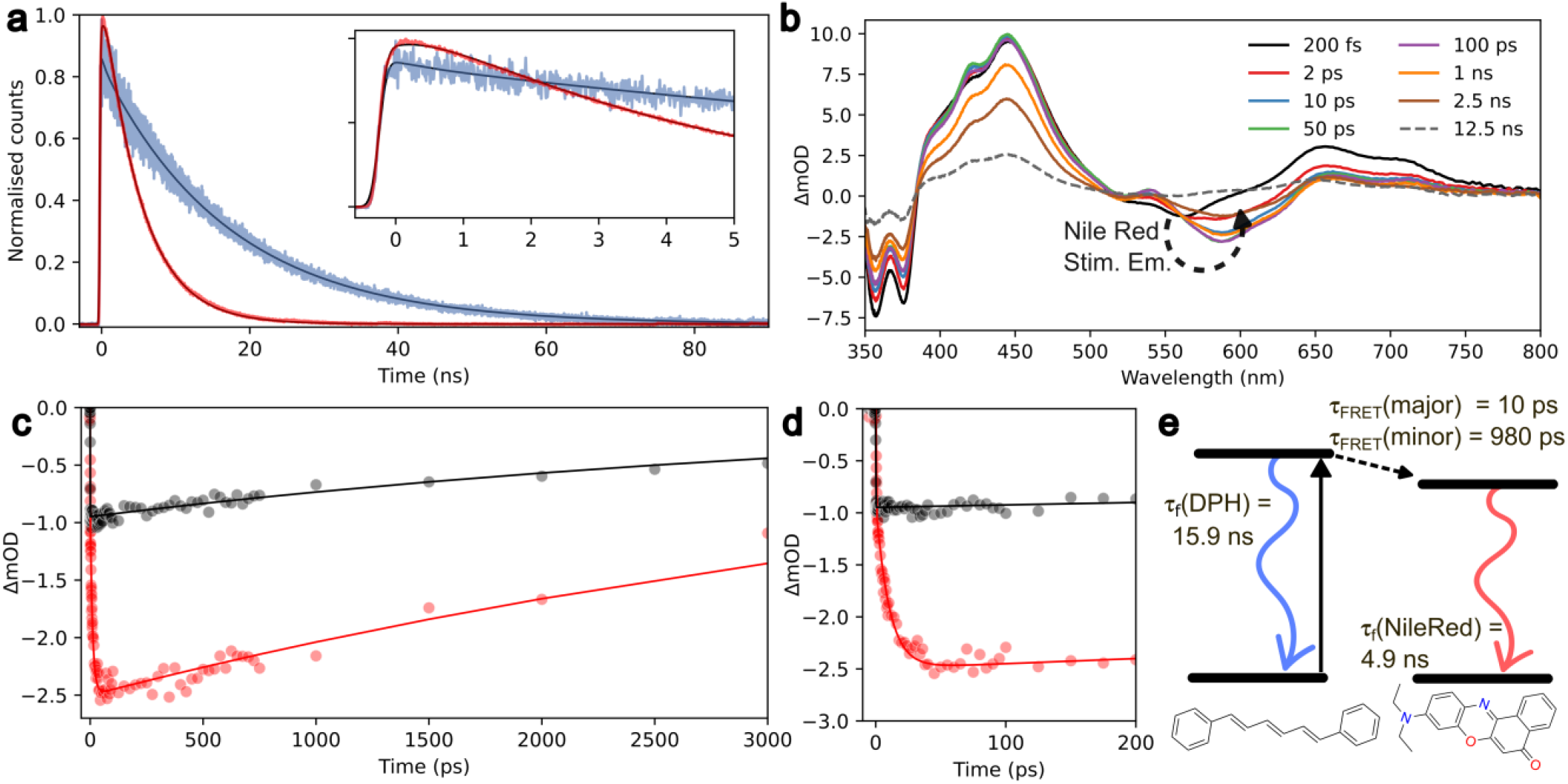
TCSPC and transient absorption measurements reveal ultrafast FRET between DPH and Nile Red. (a) Fluorescence decay in the heptamer for DPH (blue) and Nile Red (red) after excitation at 352 nm. (b) Wavelength resolved transient absorption of the heptamer for DPH and Nile Red after 352 nm excitation for 8 different pump-probe time delays. ΔmOD: change in milli-optical density. (c) Kinetics obtained by integration over the Nile Red stimulated emission signal for the heptamer with Nile Red after 535 nm excitation (black circles) and the heptamer with DPH and Nile Red using 352 nm photoexcitation (red circles), and overlaid fits (solid lines). (d) Early time dynamics of the data shown in (c) illustrating the difference between photoexcitation of DPH or Nile Red on the Nile Red stimulation emission kinetics. (e) Kinetic model of FRET and radiative decay pathways determined for DPH and Nile Red FRET in αHBs. TCSPC conditions: 3 μM DPH and Nile Red, 5 μM peptide assembly, HEPES, 10% v/v MeCN, pH 7. TA conditions: 10 μM DPH and Nile Red, 15 μM peptide assembly, HEPES, 10% v/v MeCN, pH 7. For TCSPC fitting parameters and time-resolved emission spectra, see Tables S4 – S7 and Figures S17-S18. For TA fitting parameters and wavelength resolved transient absorption, see Table S8 and Figure S20.

The relative amplitude of the fluorescence rise component in the TCSPC measurements was surprisingly small despite sizeable cross-peaks in 2D fluorescence maps and negligible direct excitation of Nile Red (Figures 2 and S6). This implies that most Nile Red molecules undergo energy transfer on timescales faster than the instrument response function (IRF) for the TCSPC experiment (170 ps).

To probe the possibility of faster FRET dynamics in αHBs, transient absorption (TA) spectroscopy with three orders of magnitude higher time resolution (IRF ∼280 fs), was applied to the heptameric and octameric barrels (Figure 3b-d). Spectral assignments of the main features of the TA data are given in the SI (Figure S20). Critically, a negative signal centered at 590 nm rises within the first few picoseconds associated with stimulated emission from Nile Red in both heptamer and octamer barrels (see kinetics in Figure 3c,d). The delayed rise originates from prompt energy transfer between DPH and Nile Red, and subsequent fluorescence from Nile Red. In contrast, control measurements that directly excited Nile Red (Figure S20) showed this band immediately (Figures 3c,d) confirming the delayed rise must be from FRET. Analysis of the data returned fast FRET time constants between DPH and Nile Red: 10.4 ± 2.4 ps (Figure 3c, Table S8) in the heptamer, and 7.9 ± 5.7 ps (Figure S22, Table S8) in the octamer. This prompt FRET timescale translates to predicted inter-chromophore separations of 1.37 ± 0.05 and 1.24 ± 0.15 nm (Figure S23), respectively, reconciling the high FRET efficiency determined from steady state emission measurements. A secondary ∼1 ns rise in the stimulated emission signal was not evident within the TA data, as would be expected from TCSPC measurements. We suggest that this is because the ∼1 ns FRET component corresponds to a minority population which is only observed in TCSPC measurements due to the technique’s sensitivity to only fluorescence and higher signal-to-noise ratios.

We rationalize these different observations by the presence of two conformationally different states present in both of the investigated αHBs: 1) a dominant population where DPH and Nile Red are in close proximity facilitating rapid energy transfer on a ∼10 ps timescale; and 2) a minor population where DPH and Nile Red are located ∼3 nm apart within a barrel and undergo a slower energy transfer with a time constant of ∼1 ns (Figure 3e).

### 2.3 MOLECULAR DYNAMICS SIMULATIONS INDICATE FAVOURABLE STACKING BETWEEN DYES

To investigate the two states proposed from ultrafast spectroscopy measurements further, we used docking and molecular dynamics (MD) simulations to explore the DPH and Nile Red binding conformations within the αHBs. Three 0.5 μs MD simulations of a DPH:Nile Red:heptamer complex were initiated from three distinct poses. The latter were generated by simultaneously docking DPH and Nile Red into the αHB using AutoDock4 Vina (Figure S24).^46^ The MD simulations indicated that the complexes were stable, with DPH and Nile Red remaining either fully π-stacked within 3.5 Å or slipped-stacked within 10.0 Å distances (measured from their centers of mass) throughout simulations in the heptamer (Figure S25). The overall αHB conformation did not change significantly from the determined X-ray crystal structure through the trajectories, with the majority of C-alpha backbone root mean square fluctuation ≤0.5 Å (Figures S26 - S29). Similar results were obtained for the octamer αHB (Figures S30 – S35).

Since the ligand movement along the channel (z) axis was slow (Figures S28 and S34), metadynamics simulations were initiated for the three docking poses within the heptamer to ensure that local minima of the energy landscape were explored. The distance between DPH and Nile Red was used as a bias variable and sampled from 1.5 Å to 40 Å to observe many conformations along the entire channel. After 1 μs, all three simulations converged to the same free-energy surface irrespective of the starting position of the simulation (Figure 4a). The free-energy surface had 2 deep minima at 3.5 Å and 10.0 Å corresponding to the distances observed with docking and traditional MD simulations (Figures 4a and S24).

**Figure 4.**
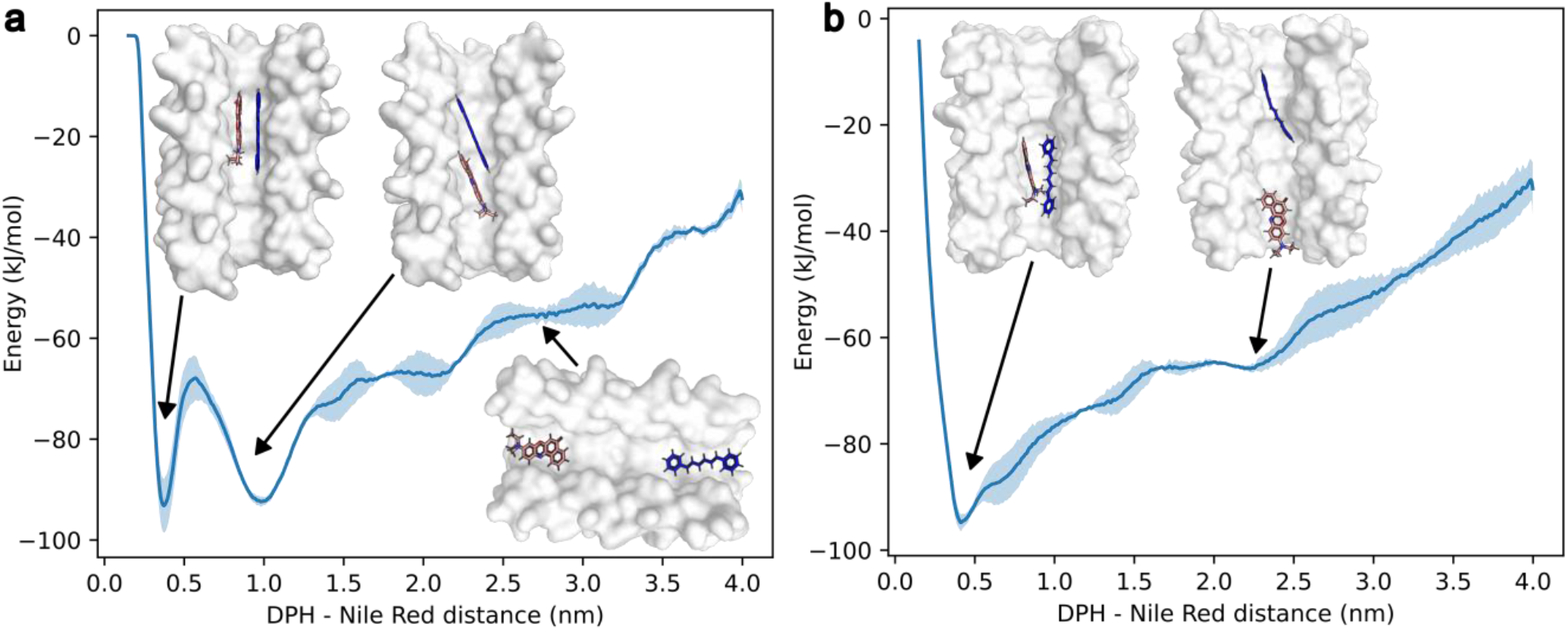
Free energy landscape sampling the distance between DPH (blue) and Nile Red (red) within the αHBs. (a) The heptamer has two minima at 3.5 Å (–93 ± 5 kJ mol^-1^) and 10.0 Å (–92 ± 1 kJ mol^-1^). (b) The octamer shows a broad global minimum at 4.1 Å (–94.6 ± 2 kJ mol^-1^). 3 × 1 μs simulations were started independently from the starting poses in Figures S24 and S30. Standard deviation from the mean is shown in pale blue.

Shallower local minima were also observed between 20 – 30 Å. For the octamer, we observed a broad global minimum at 4.1 Å and a shallower minimum around 20 Å (Figure 4b). The closer internuclear distance between DPH and Nile Red matched well with the ∼10 ps ultrafast energy transfer time constant between the dyes observed in TA spectroscopy. The shallower higher energy minima at greater dye separations will have a lower population at room temperature, and thus represents a small percentage of the total sample, and this possibly corresponds to the slower ∼1 ns dynamics extracted with TCSPC measurements.

### 2.4 αHBS ENCAPSULATE ANTHRACENE AND CATALYSE ITS PHOTODIMERISATION

Encouraged by the successful demonstration of FRET between DPH and Nile Red with clear co-occupancy of the dyes in the heptameric and octameric barrels, and the modelling indicating intermolecular π-stacking (Figure 4), we investigated if similar interactions could be promoted and exploited in αHBs for different hydrophobic molecules. Anthracene was chosen as, when pre-organized into a π-stacked dimer, it undergoes a well-characterized [4+4] cycloaddition reaction upon UV-irradiation at 365 nm. This is readily followed spectroscopically even at μM concentrations, as photodimerization reduces anthracene emission.

*In silico* docking of two anthracene molecules into the heptamer predicted two poses: a potentially reactive π-stacked dimer with a 3.5 Å separation, and a slip-stacked conformation (Figure 5a). Starting from these poses, 0.5 μs MD simulations revealed the ternary complex to be highly dynamic, with anthracene molecules in an equilibrium between a monomer and a π-stacked dimer (Figure S36). By contrast, docking with the hexamer indicated that its diameter would be insufficient to accommodate two fully π-stacked anthracene molecules (Figure S37). We confirmed anthracene encapsulation within both αHBs experimentally through co-sedimentation during ultracentrifugation (Figure S38), and by observing an induced circular dichroism (CD) signal (Figure 5b). The latter is a consequence of the Cotton effect, and only evident when anthracene molecules are encapsulated in a chiral environment, *i*.*e*., like the lumen of an αHB.^47^ As before, we used the trimeric peptide as a control. For the trimer, anthracene co-sedimentation was observed, but no induced CD signal was evident, confirming only non-specific interactions between anthracene and the trimer peptide (Figures S38 and 5b).

**Figure 5.**
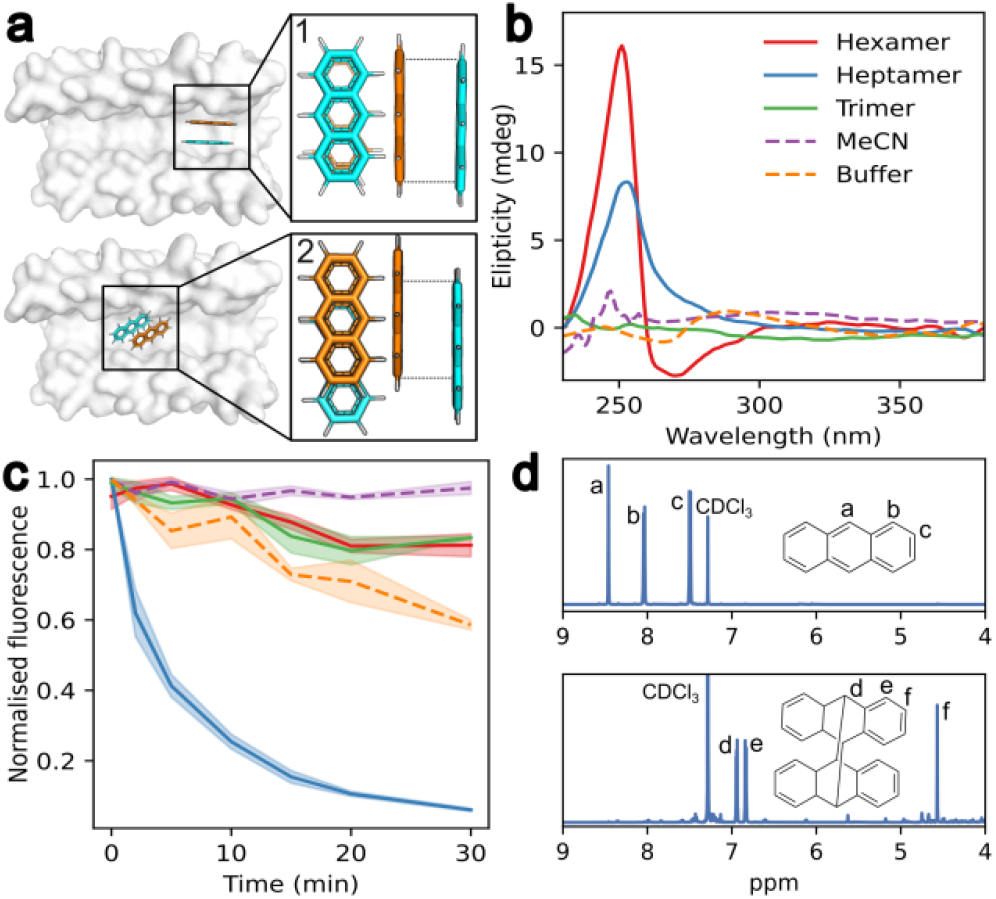
αHB channel dimensions control anthracene photodimerization in aqueous solution. (a) Lowest energy AutoDock Vina poses for anthracene dimer in the heptamer, showing reactive (1) and offset (2) conformations. (b) Induced CD signal upon anthracene binding to the heptamer and the hexamer. (c) Decrease in anthracene emission upon irradiation at 365 nm. Color key same as in (b): hexamer, red; heptamer, blue; trimer, green; MeCN, purple; buffer, orange. (d) Proton NMR spectra without illumination (top) and after 30 mins of excitation at 365 nm (bottom) for anthracene dimerization reaction within the heptamer. Conditions: 14 μM anthracene, 7 μM peptide assembly, HEPES, 10% v/v MeCN, pH 7, and peptides were removed before collecting NMR data.

Based on modelling and the CD data, we hypothesized that the heptamer could catalyze anthracene dimerization while the hexamer should inhibit it. When mixed in a 1:2 (αHB:anthracene) ratio and irradiated with 365 nm light, the photodimerization reaction proceeded to completion with the heptamer as indicated by a decrease in fluorescence signal over 30 minutes (Figure 5c). Complete conversion to the photodimer product was confirmed by ^1^H NMR, (Figure 5d). By contrast, no clear reaction was observed with the hexamer (Figure 5c), other than a much slower, linear loss of anthracene fluorescence similar to buffer-only, acetonitrile, and trimer controls. These experiments show how encapsulation of the anthracene within the heptamer greatly accelerates photodimerization, presumably by binding two anthracene molecules in a complex primed for [4+4] cycloaddition. Moreover, it highlights that not all αHBs are compatible with the transformation, and that selectivity can be achieved by appropriate choice of barrel, which can be predicted by modelling.

## 3. CONCLUSIONS

Here we demonstrate the ability to bind and organize two ligands in the lumens of *de novo* designed α-helical barrel assemblies (αHBs) in conformations that accelerate the timescales for two different photoinduced chemical processes. First, we show that by exploiting different sizes and shapes of their hydrophobic lumens, we can select αHBs that facilitate energy transfer between two fluorophores, DPH and Nile Red. Using ultrafast spectroscopy, we reveal that the channels of heptameric and octameric peptide assemblies are wide enough to encapsulate both dyes simultaneously and juxtapose them for efficient energy transfer on picosecond timescales. Molecular dynamics simulations predict that DPH and Nile Red bind in π-stacked complexes facilitating the observed fast energy transfer. Moreover, by varying the internal diameter of the structures (∼7 Å in the hexamer *vs*. ∼8 Å in the heptamer), we can select for barrels that instead of promoting FRET, preferentially encapsulate only one of the dyes. These two features – ligand encapsulation with (i) selective binding and (ii) in privileged conformations for catalysis – are the key features of enzymes. Here, we achieve these with shape complementarity and non-specific hydrophobic interactions within symmetric peptide assemblies as compared to the more-complex active sites found in natural enzymes. To illustrate the potential of the αHBs in catalysis in aqueous buffers, we explore the generation of tightly packed anthracene dimers that are primed for a photodimerization reaction. We find that cycloaddition can be catalyzed within a heptameric αHB, but not within the narrower hexamer. Again, as indicated by predictive modelling, these behaviors appear to be controlled by the relative sizes and shapes of internal lumens of the αHBs.

In the broader context of *de novo* protein design, current approaches towards developing small-molecule binders and catalysts often rely on deep-learning methods, which do not necessarily enhance our understanding of protein function. In contrast, we control binding and catalysis in a rational and predictive manner by leveraging the concept of “molecular flasks” borrowed from supramolecular chemistry.

Looking ahead, these peptide-based *de novo* designed αHBs are thermostable and tolerate a large number of mutations in their lumens,^41^ including to polar, charged, aromatic and non-canonical side chains. As a result, a wide range of small molecules could be encapsulated within the αHBs for manipulation.^41^ Moreover, recently, we have shown that the αHB peptide assemblies can be converted to thermostable single-chain proteins by linking multiple helices together through computational protein design.^48^ These single-chain proteins are produced by expression from synthetic genes, which opens possibilities for desymmetrization and functionalization through further rational computational protein design, and to improve activity using directed evolution.^6, 49^ Thus, the αHB peptides and proteins offer an exciting platform for combining the shape complementarity and confinement offered by organic molecular flasks with the diversity of binding-site geometries and substrate selectivity seen in natural enzymes.

## ASSOCIATED CONTENT

### Supporting Information

Experimental materials and methods, peptide sequences, additional steady state and ultrafast spectroscopy data and fits, additional docking and molecular-dynamics simulations data (PDF).

The raw data and code used in this publication have been deposited at Zenodo (DOI: 10.5281/zenodo.13335904).

## AUTHOR INFORMATION

### Author Contributions

RP, DAS, TAAO and DNW conceived the study and contributed to the experimental design. RP designed and screened the peptides. RP solved the 9f5a crystal structure. RP and OAH collected the TCSPC data. RP and JFJ collected the TA data. JMF, UO, LL and FP contributed to peptide characterization. RP, DAS, TAAO and DNW wrote the manuscript with contributions from all authors. DNW, DAS, GJL and TAAO provided mentorship to RP, OAH, JFJ, FP, LL, JMF and UO.

## Notes

RP is a South West Biosciences Collaborative Awards in Science and Engineering student supported by Rosa Biotech. DAS, JMF and UO were employees at Rosa Biotech from 2019 to 2024.

DNW is a cofounder and director of Rosa Biotech. All other authors have no competing interests.

## ACKNOWLEDGMENTS

RP is supported by a BBSRC-funded PhD studentship and by Rosa Biotech (South West Biosciences Doctoral Training Partnership). FP was supported by an Engineering and Physical Sciences Research Council (EPSRC) program grant to GJG and DNW (EP/T012455/1). OAH, JFJ and TAAO acknowledge financial support from EPSRC for the award of Programme Grant EP/V026690/1. TAAO acknowledges support from the Royal Society for a University Research Fellowship (UF1402310, URF\R\201007) and Research Fellows Enhancement Award (RF\ERE\210045). We thank the Mass Spectrometry Facility, School of Chemistry, University of Bristol, for access to the EPSRC-funded Bruker Ultraflex MALDI–TOF instrument (EP/K03927X/1). We would like to thank Diamond Light Source for access to beamline I04 (proposals mx23269 and mx31440). We thank internship students at Rosa Biotech, Josh Lewin and Dr Nokomis Ramos-Gonzalez, who conducted ligand docking and molecular dynamics simulations of multiple dyes in αHB, that led to the current work. Finally, we thank the members of the Woolfson and Oliver laboratories for many helpful discussions.

## ABBREVIATIONS

αHB: alpha-helical barrel
CD: circular dichroism
DPH: 1,6-diphenyl-1,3,5-hexatriene
FRET: Förster resonance energy transfer
MD: molecular dynamics
TA: transient absorption
TCSPC: time-correlated single photon counting.

## Notes

### Competing Interest Statement

The authors have declared no competing interest.

