## Supporting Information for "Confinement and Catalysis Within *De Novo* Designed Peptide Barrels"

### 1 Methods

#### 1.1 General

All solvents, chemicals, and reagents were purchased from commercial sources and used without further purification. Fluorenylmethoxycarbonyl(Fmoc)-α-L-amino acids, Rink amide MBHA resin for solid-phase peptide synthesis (SPPS) and N,N-dimethylformamide (DMF) were purchased from Sigma-Aldrich and Cambridge Reagents. Coupling reagents Oxyma Pure and diisopropylcarbodiimide (DIC) were purchased from Fluorochem. Morpholine, trifluoroacetic acid (TFA) were purchased from Sigma-Aldrich; pyridine was from Thermo Fisher; triisopropylsilane (TIPS) was from Acros Organics. All other chemicals were reagent grade and purchased from Sigma-Aldrich. Peptide biophysical characterization were recorded in HEPES buffer (25mM HEPES, 100mM NaCl, pH 7). Peptide characterisation data for CC-Type2-[$S_gL_aI_d$], CC-Type2-[$I_aV_d$], CC-Type2-[$S_gI_aI_d$], CC-Type2-[$V_aV_d$], CC-Type2-[$L_aI_d$], CC-Type2-[$L_aI_d$]-L28Y, CC-Type2-[$L_aI_d$]-L14A, CC-Type2-[$L_aI_d$]-I24A, CC-Type2-[$L_aI_d$]-I17H, CC-Type2-[$L_aI_d$]-L21S-I24Y, CC-Type2-[$L_aI_d$]-I24Y, CC-Type2-[$L_aI_d$]-L7Y, CC-Type2-[$L_aI_d$]-L21K and the trimer has been previously published. [1,2]

#### 1.2 Computational tools

##### 1.2.1 ISAMBARD

$\alpha$HBs for the octamer with the oligomer state of 7 and 8 were modelled using the in-build GA optimizer and the BUFF force-field with a population size of 200 and the generation number of 10.[3]

##### 1.2.2 AutoDock Vina

3D ligand structures for DPH, Nile Red and anthracene were taken from PubChem database, geometry-optimised with MOPAC (PM6-D3H4 Hamiltonian with COSMO solvation) and docked with AutoDock Vina 1.2 with exhaustiveness of 64.[4,5] Vina input files were generated using Babel; the number of DPH torsion angles was set to 0.[6]

##### 1.2.3 Molecular dynamics simulations

OpenMM 7.7 was used for running MD simulations.[7] All simulations were run using a Langevin integrator with an integration step of 2 fs and a collision rate of 1 $ps^{-1}$, the pressure was kept constant at 1.01325 bar with a MonteCarlo barostat. All systems were solvated with tip3p waters and 100 mM NaCl in a rhombic dodecahedron with a padding of 1 nm. Particle mesh Ewald with a cut-off of 1 nm was used for long-range electrostatics. Simulations were run with the Amber ff14SB force field; small molecule parameters were generated using the SMIRNOFF force field. The initial peptide-ligand complexes were prepared by inspecting the top 25 DPH and Nile Red poses from AutoDock Vina[4], and taking the lowest energy pose for each conformation.

The peptide-ligand complex was initialised at 10 K with 5 kcal/mol restraints on the peptide backbone and ligand heavy atoms and minimised. The system was then heated to 293 K over 600 ps. Heavy atom restraints were relaxed over 3.5 ns and the unrestrained system was further equilibrated for 10 ns. A production run was then performed for 500 ns. Periodic boundary conditions were removed using GROMACS trajconv[8], trajectories were analysed using MDAnalysis.[9]

For metadynamics, equilibrated systems were run without the MonteCarlo barostat for 1000 ns with an initial energy barrier of 0.5 kJ/mol and a frequency of 5000 steps. The distance between the centres of mass of DPH and Nile Red was used as the bias variable and was sampled between 0.15 and 4 nm with a step of 0.01 nm.

#### 1.3 Peptide synthesis and purification

##### 1.3.1 Automated microwave Fmoc/tBu solid-phase peptide synthesis (SPPS)

Automated microwave SPPS was performed on a Liberty Blue (CEM) synthesizer with inline UV monitoring. Syntheses were performed on 0.1 mmol scales with side-chain protection of the amino acids as follows: Gln(Trt), Glu(O*t*Bu), Lys(Boc), Tyr(*t*Bu), Trp(Boc). The resin (Rink amide MBHA, 0.65 mmol/g loading, 100 – 200 mesh) was weighted to enable a synthesis on a 0.1 mmol scale. The coupling reactions were performed by adding protected amino acids dissolved in DMF (2.5 mL, 0.2 M), the coupling reagent DIC in DMF (1.0 mL, 1 M) and Oxyma Pure in DMF (1 mL, 0.5 M) to the respective resin. Standard couplings were performed at 90 °C for 4 min (100 W for 20 s, 60 W for 10 s, 35 W for 240 s). Standard deprotections were performed using 20% (v/v) morpholine in DMF at 90 °C for 1 min (125 W 30 s, 32 W 60 s). All peptides were manually acetyl capped through addition of pyridine (0.5 mL) and acetic anhydride (0.25 mL) in DMF (9.25 mL), shaking at room temperature (rt) for 20 minutes. The resin was washed three time with DMF followed by six times with DCM before cleavage. Peptides were cleaved from the resin with addition of 10 mL of a mixture 95:2.5:2.5 v/v trifluoroacetic acid (TFA)/$H_2O$/triisopropylsilane (TIPS), shaking at room temperature for 2 hours. The TFA solution was then filtered to remove the resin beads and was reduced in volume to ≈5 mL or lower using a flow of $N_2$. Cleaved peptide was precipitated with cold diethyl ether (≈45 mL), isolated via centrifugation and dissolved in a 1:1 mixture MeCN/$H_2O$. Crude peptides were lyophilized to yield a white or off-white powder.

##### 1.3.2 Semi-preparative High Performance Liquid Chromatography (HPLC)

All peptides were purified by reverse phase HPLC (JASCO) using a Luna C18 (Phenomenex) column (150 x 10 mm, 5 μm particle size, 100 Å pore size). Crude peptide was dissolved at 7 mg/mL in 40% v/v MeCN in $H_2O$ with 0.1% TFA, injected to the column and eluted with a 3 mL/min linear gradient (40 – 100%) of MeCN in $H_2O$ with 0.1% TFA each over 30 minutes. Elution of the peptide was detected with inline UV monitoring at 220 and 280 nm wavelengths simultaneously. A column oven (50 °C) was employed to improve separation. Pure fractions were identified by analytical HPLC and matrix-assisted laser desorption/ionization–time of flight (MALDI-TOF) mass spectrometry, then pooled, and freeze-dried.

##### 1.3.3 Analytical HPLC

Analytical HPLC traces were obtained using a Jasco 2000 series HPLC system and a Phenomenex Kinetex C18 (100 x 4.6 mm, 5 μm particle size, 100 Å pore size) column. Chromatograms were monitored at 220 and 280 nm wavelengths. The linear gradient was 40 – 100% MeCN in water (each containing 0.1% TFA) over 25 min at a flow rate of 1 mL/min. When required, a column oven (50 °C) was employed to assist peptide elution.

##### 1.3.4 Mass spectrometry

Matrix-assisted laser desorption/ionization–time of flight (MALDI-TOF) mass spectra were collected on a Bruker UltraFlex MALDI-TOF mass spectrometer operating in positive-ion reflector mode. Peptides were spotted on a ground steel target plate using α-cyano-4-hydroxycinnamic acid dissolved in 1:1 MeCN/$H_2O$ as the matrix. Masses quoted are for the monoisotopic mass as the singly protonated species.

#### 1.4 Solution-phase biophysical characterizations

##### 1.4.1 Peptide and ligand concentration determination

Peptide concentration was determined at 280 nm using a Nanodrop 2000 (ThermoScientific) spectrometer ($\varepsilon_{280}$ = 5690 $M^{-1}cm^{-1}$). DPH ($\varepsilon_{356}$ = 84000 $M^{-1}cm^{-1}$), Nile Red ($\varepsilon_{535}$ = 48000 $M^{-1}cm^{-1}$), Coumarin-7 ($\varepsilon_{426}$ = 52500 $M^{-1}cm^{-1}$), Methyl orange ($\varepsilon_{466}$ = 26000 $M^{-1}cm^{-1}$) and anthracene ($\varepsilon_{356}$ = 9700 $M^{-1}cm^{-1}$) concentrations were determined using Cary 100 UV-Vis (Agilent Technologies).

##### 1.4.2 Circular dichroism (CD) spectroscopy

Circular dichroism (CD) data were collected on a JASCO J-810 or J-815 spectropolarimeter fitted with a Peltier temperature controller. Peptide samples were acquired at 50 μM at 5 °C in HEPES buffer. Data were collected in a 1 mm quartz cuvette between 190 and 260 nm with the instrument set as follows: band width 1 nm, data pitch 1 nm, scanning speed 100 nm/min, 1 s response time. Each CD spectrum was obtained by averaging of 8 scans.

Anthracene binding data was acquired at 15 μM peptide assembly, 30 μM anthracene at 5 °C in HEPES buffer with 10% MeCN. Data were collected in a 1 cm quartz cuvette between 240 and 300 nm with the instrument set as follows: band width 1 nm, data pitch 1 nm, scanning speed 500 nm/min, 1 s response time. Each CD spectrum was obtained by averaging of 10 scans. CD spectra of 15 μM peptide assembly under the same conditions was used as a blank.

For peptide thermal-response experiments, the CD signal at 222 nm wavelength was monitored over the temperature range 5 – 95 °C at a ramp rate of 60 °C per hour and with the same settings and peptide or protein concentrations given above.

Where applicable, the spectra were converted from ellipticities (mdeg) to mean residue ellipticities (MRE, (deg $cm^2$ $dmol^{-1}$ $res^{-1}$)) by normalizing for concentration of peptide bonds and the cell path length using the equation:

$$(1) \qquad MRE = \frac{\theta \times 10^6}{c \times l \times n}$$

where the variable $\theta$ is the measured difference in absorbed circularly polarized light in millidegrees, $c$ is the µM concentration of the compound, $l$ is the path length of the cuvette in mm, and $n$ is the number of amide bonds in the polypeptide.

##### 1.4.3 Analytical ultracentrifugation

Analytical ultracentrifugation (AUC) was performed on a Beckman Optima X-LA or X-LI analytical ultracentrifuge with an An-60-Ti rotor (Beckman-Coulter). Buffer densities, viscosities, and peptide partial specific volumes ($\overline{v}$) were calculated using SEDNTERP (http://rasmb.org/sednterp/).

For sedimentation velocity (SV), peptide sample solutions of 310 or 410 µL were prepared in HEPES at 150 µM peptide concentration and placed in a sedimentation velocity cell with an epon or aluminium, respectively, 2-channel centrepiece and quartz windows. The reference channel was loaded with 320 µL (or 420 µL, respectively) of HEPES buffer. The samples were centrifuged at 50 krpm at 20 °C, with absorbance scans taken over a radial range of 5.8 – 7.3 cm at 5 min intervals to a total of 120 scans. Data from a single run were fitted to a continuous c(s) distribution model using SEDFIT[10] at 95% confidence level.

For anthracene co-sedimentation, 15 µM peptide assembly with 15 µM anthracene was prepared in HEPES, 10% v/v MeCN. The sample was placed in a sedimentation velocity cell with an aluminium 2-channel centrepiece and centrifuged at 50 krpm at 20 °C, with absorbance scans taken at 250 nm over a radial range of 5.8 – 7.3 cm at 5 min intervals to a total of 120 scans. Data from a single run were fitted to a continuous c(s) distribution model using the peptide partial specific volumes or to a 'bias-free' g*(s) distribution using SEDFIT as described above.

Sedimentation equilibrium experiments were performed at 70 µM peptide concentration in 110 µL at 20 °C. The experiment was run in triplicate in a six- channel centrepiece. The samples were centrifuged at speeds in the range of or 20 – 45 krpm and scans at each recorded speed were duplicated after equilibration for 8 hours. Data were fitted using SEDPHAT[11] to a single species model. Monte Carlo analysis was performed to give 95% confidence limits.

##### 1.4.4 Ligand binding

DPH and Nile Red binding experiments were pipetted in quadruplicate using an epMotion 5070 liquid handler (Eppendorf). The total concentration of ligand was kept constant (0.5 µM in 10% v/v MeCN) and the concentration of *de novo* peptide assembly varied from 0 – 30 µM. Data were collected on a Clariostar plate reader (BMG Labtech) (DPH $\lambda_{ex}$ = 350 ± 8 nm, $\lambda_{em}$ = 450 nm and Nile Red $\lambda_{ex}$ = 535 ± 8 nm, $\lambda_{em}$ = 593 nm). Binding constants were extracted by fitting the data to the following equation:

$$(2) \qquad y = B_{max} \frac{(c + x + K_D) + \sqrt{(c + x + K_D)^2 - 4cx}}{2c}$$

where *c* is the total concentration of the constant component (*e.g.* DPH), *x* is the concentration of variable component (*e.g.* peptide/protein), $B_{max}$ is the fluorescence signal when all of the constant component is bound, and *y* is the fraction of bound component being monitored *via* fluorescence signal.

##### 1.4.5 FRET screen and steady state spectra

Coumarin-7, Methyl orange, DPH and Nile Red fluorescence data was collected on a Clariostar plate reader (BMG Labtech) at 5µM of the *de novo* peptide assembly, 3 µM ligand in HEPES, 10% v/v MeCN. DPH $\lambda_{ex}$ = 350 ± 8 nm, Nile Red $\lambda_{ex}$ = 535 ± 8 nm, gain was kept below 2500.

2D excitation-emission measurements for the FRET screen hits were performed on a FS5 spectrofluorometer (Edinburgh Instruments) with $\lambda_{ex}$ from 280 nm to 600 nm, $\lambda_{em}$ from 350 nm to 700 nm with a dwell time of 0.25 s and 2 nm bandwidth.

##### 1.4.6 Anthracene photodimerization

The reaction was performed by illuminating the samples with 365 nm light (M365LP1, Thorlabs) at 14 µM anthracene, 7 µM peptide assembly in HEPES, 10% v/v MeCN, pH 7. The reaction was followed by diluting 10 µL of the sample into 90 µl MeCN and measuring the fluorescence after excitation at 365 nm with the Clariostar plate reader (BMG Labtech). This reduced the noise from scattering and crashing-out of anthracene in buffer and ccTri samples.

NMR samples were prepared from a 10ml reaction. Peptide was precipitated with $CHCl_3$ and isolated via centrifugation. Remaining $CHCl_3$ was evaporated, and the products were redissolved in 40 µl $CDCl_3$. $^1$H NMR spectra was collected on Bruker Avance III HD 700. All chemical shifts are quoted in parts per million relative to tetramethylsilane (0.00 ppm) and coupling constants are measured in Hz.

Before irradiation (anthracene): $^1$H NMR (700 MHz, $CDCl_3$) δ 8.46 (s, 1H), 8.04 (dt, *J* = 6.0, 3.0 Hz, 2H), 7.49 (dt, *J* = 6.0, 3.0 Hz, 2H).

After irradiation (anthracene dimer): $^1$H NMR (700 MHz, $CDCl_3$) δ 6.94 (dd, *J* = 5.4, 3.3 Hz, 1H), 6.84 (dd, *J* = 5.5, 3.1 Hz, 1H), 4.57 (s, 1H). Impurities due to peptide contamination: 2.38 (t, *J* = 7.5 Hz, 0H), 1.70 (s, 1H), 1.52 (s, 2H), 1.45 (d, *J* = 10.3 Hz, 1H), 1.39 – 1.34 (m, 1H), 1.32 (d, *J* = 4.4 Hz, 1H), 1.31 (s, 4H), 1.28 (s, 7H), 1.26 (s, 1H), 1.23 (s, 1H), 1.10 (s, 5H), 0.90 (dd, *J* = 8.3, 5.7 Hz, 2H), 0.90 – 0.80 (m, 14H).

#### 1.5 Ultrafast Laser Spectroscopy

##### 1.5.1 Time-Correlated Single Photon Counting

Time-correlated single photon counting (TCSPC) data were acquired using a home-built TCSPC apparatus previously reported.[12] The output of a tunable high-power ultrafast oscillator (Chameleon Ultra II, 3.7 W, 80 MHz, Coherent) was frequency doubled to generate the required 352 nm pulses for DPH excitation. For Nile Red, the light was first frequently shifted to 1070 nm with an optical parametric oscillator (Chameleon Compact, APE) and then frequency doubled to generate 535 nm excitation pulses. To avoid re-excitation of samples, the repetition rate was reduced to 6.67 MHz using a pulse picker (cavity dumper, APE). The diffracted output was spatially filtered with a pinhole to remove residual zeroth order diffraction, and the resulting light was focused into a stirred liquid sample (1 cm path length cell). Fluorescence was collected at 90° relative to excitation using an infinity-corrected microscope objective (4×/0.2 NA Plan Apochromat, Nikon) and through a series of achromatic lenses collimated and focused onto an avalanche photodiode detector (ID100-50-ULN, IDQ).

Photon count arrival times were recorded with a time-to-digital converter (Time Tagger 20, Swabian Instruments) and accumulated in 10 ps bins. To facilitate wavelength-resolved TCSPC measurements, fluorescence was propagated through a birefringent interferometer (Gemini, Nireos) after the collection objective and before the detector.[13] Each data point was integrated for 1 s, and every spectral interferogram was averaged 10 times. All TCSPC data were acquired by using customized LabVIEW software (National Instruments). The wavelength-resolved data was not corrected for spectral filtering due to transmission through the collecting objective and Gemini interferometer.

For the weakly fluorescing hexamer sample with DPH and Nile Red, Nile Red fluorescence was collected by replacing the birefringent Gemini interferometer with a >565 nm filter.

The instrument response function (IRF) was 170 ps, as determined by recording laser scatter from solvent in a 1 cm path length cuvette. All data were collected at room temperature (20 °C) using the magic angle condition at 5μM of the *de novo* peptide assembly, 3 μM ligand in HEPES, 10% v/v MeCN, pH 7.

##### 1.5.2 Transient Absorption

A synchronised dual amplifier laser system (Pharos PH2, Light Conversion) was used for transient absorption (TA) experiments. The first amplifier was operated at 0.5 kHz and used to pump an optical parametric amplifier (Orpheus-HP, Light Conversion) to generate excitation pulses which were tuned to excite either DPH (352 nm) or Nile Red (535 nm). A small fraction of the 1030 nm fundamental output from the second amplifier (operating at 1 kHz) was focused into a continually rastered $CaF_2$ window to generate a white light supercontinuum probe spanning 350–800 nm. Residual fundamental 1030 nm was removed from the supercontinuum using a Schott glass filter. The pump-probe time delay was controlled by a mechanical delay stage and used to generate delays between –10 ps to +13 ns. Pump and probe laser beams were focused into a 1 mm pathlength flow cell (Starna), and the transient signal and colinear probe beam were collimated and imaged into a homebuilt spectrograph coupled to a CCD linear array detector (Stresing). Pump and probe pulses were polarized at magic angle with respect to each other. The sample absorbance for TA experiments was ~ 0.03 OD (1 mm pathlength) at 352 nm, and the solutions were circulated throughout experiments. The instrument response function was 280 fs as measured *via* non-resonant buffer response.

##### 1.5.3 DPH fluorescence quantum yield determination

DPH quantum yields within the αHB were estimated using DPH quantum yield in acetonitrile ($\Phi_{MeCN}$ = 0.19) as a reference.

(3) $\phi_{aHB} = \phi_{MeCN}\ x\ \frac{n^2}{n_{MeCN}^2}\ x\ \frac{F_{aHB}}{F_{MeCN}}$, where $n$ is the refractive index of the buffer (1.33) and $n_{MeCN}$ is the refractive index of acetonitrile (1.34).

The absorbance was varied from 0.01 to 0.08, peptide concentration was kept fixed at 5 μM of the *de novo* peptide assembly in HEPES, 10% v/v MeCN, pH 7. Measurements were done in triplicate and averaged.

##### 1.5.4 Kinetic fits, Global analysis and calculation of FRET distance

Lifetime decays after the direct excitation of DPH and Nile Red were modelled as a mono- or bi-exponential decay convolved with a Gaussian IRF function. See Tables S4-S7.

The energy transfer from the donor (DPH) was modelled as a bi-exponential decay accounting for the population of DPH ($A_2$) that are not involved in FRET:

$$(4) \qquad I_{DPH} = A_1 e^{-(1/\tau_{FRET}+1/\tau_{DPH})t} + A_2 e^{-t/\tau_{DPH}}$$

The energy transfer to the acceptor (Nile Red) was modelled as a bi-exponential decay where $B_1$ is a negative amplitude reflecting the rise of acceptor fluorescence due to FRET:

$$(5) \qquad I_{NR} = B_1 e^{-t/\tau_{FRET}} + B_2 e^{-t/\tau_{NR}}$$

These were fitted to DPH and Nile Red channels simultaneously while floating the exponential pre-factors ($A_1$, $A_2$, $B_1$, $B_2$) and $k_{FRET}$. Average $k_{DPH}$ and $k_{NR}$ were taken from Tables S4 and S6, respectively, and kept fixed. These data were acquired in control measurements of the dyes alone in αHBs. For readability, equations shown above exclude the convolution function.

Assuming $1/\tau_{FRET} = k_{FRET}$, the Förster distance was calculated as:

$$(6) \qquad R^6 = 8.785 \, x \, 10^{-5} \frac{\phi_{DPH}}{\tau_{DPH}} \frac{1}{k_{FRET}} \frac{\kappa^2 I}{n^4}$$

where $\Phi_{DPH}$ and $\tau_{DPH}$ is the fluorescence quantum yield and the lifetime of DPH inside the αHB, $\kappa$ is the orientation factor between the dipoles and $\kappa^2$ assumed to be 2/3, $I$ is the Forster spectral overlap integral of DPH fluorescence and Nile Red absorption within the αHB, and $n$ is the refractive index of the buffer (1.33). The Förster spectral overlap integral was obtained using software freely available at FluorTools.com.

In transient absorption data, the Nile Red stimulated emission (SE) signal was predominantly free from other overlapping transient features of the same chromophore or those associated with DPH in FRET experiments. Therefore, integration over feature between 560 – 600 nm probe wavelengths in Nile Red containing samples returned the kinetics associated with dye's SE. These extracted kinetics (Figure S22) were fit to a sum of exponentials convolved with a Gaussian IRF function to determine the early FRET time constants while keeping the average Nile red fluorescence decay constants to those determined in control measurements (Tables S6 and S8).

#### 1.6 Structural characterization

##### 1.6.1 Crystals growth

Diffraction-quality peptide crystals were grown using a sitting-drop vapour-diffusion method. Freeze-dried peptide was dissolved in ultrapure water and diluted to 10 mg/mL. Commercially available sparse matrix screens were used (Morpheus®, JCSG-plus™, Structure Screen 1 and 2, Pact Premier™, ProPlex™; Molecular Dimensions), and the drops were dispensed using a robot (Oryx8; Douglas Instruments). For each well of an MRC 2 drop plate, 0.3 µL of peptide solution and 0.3 µL of reservoir were mixed and the plate was incubated at 20 °C. Crystals generally formed within a month, and after looping were soaked in reservoir solution containing 25% glycerol as a cryoprotectant. Diffraction quality crystals were obtained in Pact Premier™ F11: 0.2 M sodium citrate. 0.1 M Bis Tris propane 6.5 20 % w/v PEG 3350.

##### 1.6.2 X-ray crystal structure determination

Diffraction data for the crystals were obtained at the Diamond Light Source (Didcot, UK) on beamline I04. Data were processed using the automated pipelines: Xia2 pipelines[14], which ports data through DIALS[15] to POINTLESS and AIMLESS[16] as implemented in the CCP4 suite.[17] Phases were solved by molecular replacement using the ISAMBARD model for PHASER.[18] Final structures were obtained after iterative rounds of model building with COOT[19] and refinement with Phenix Refine.[20] Solvent-exposed atoms lacking map density were either deleted or left at zero occupancy. Biological assembly was generated using PISA.[17, 21] Data collection and refinement statistics are provided in Table S2.

### 2 Supplementary Tables

#### 2.1 Biophysical and structural characterisation

**Supplementary Table 1. Sequences and associated data of all *de novo* designed aHBs used in this study.** For biophysical characterisation, see Figures S1-S2 and [1, 2]. * For simplicity, we refer to the three peptides - CC-Type2-[$S_gL_aI_d$], CC-Type2-[$I_aV_d$] and CC-Type2-[$I_aI_d$]-I17V-I21F - as the hexamer, heptamer and octamer throughout the text.

| # | Systematic name | Sequence | AUC | Crystal |
|---|---|---|---|---|
| 1 | Heptamer (CC-Type2-[$I_aV_d$]) | Ac-G EVAQAIK EVAKAIK EVAWAIK EVAQAIK G-NH2 | 6.8 | 6g66 (7) |
| 2 | Octamer (CC-Type2-[$I_aI_d$]-I17V-I21F) | Ac-G EIAQAIK EIAKAIK EVAWAFK EIAQAIK G-NH2 | 7.4 | 9f5a (8) |
| 3 | CC-Type2-[$I_aI_d$]-I17V-I21Y | Ac-G EIAQAIK EIAKAIK EVAWAYK EIAQAIK G-NH2 | 7.0 | - |
| 4 | Hexamer (CC-Type2-[$S_gL_aI_d$]) | Ac-G EIAKSLK EIAKSLK EIAWSLK EIAKSLK G-NH2 | 5.7 | 4pn9 (6) |
| 5 | CC-Type2-[$S_gI_aI_d$] | Ac-G EIAKSIK EIAKSIK EIAWILK EIAKILK G-NH2 | 6.1 | 4pnb (6) |
| 6 | CC-Type2-[$V_aV_d$] | Ac-G EVAQAVK EVAKAVK EVAWAVK EVAQAVK G-NH2 | 5.8 | 6g65 (6) |
| 7 | CC-Type2-[$L_aI_d$] | Ac-G EIAQALK EIAKALK EIAWALK EIAQALK G-NH2 | 6.3 | 4pna (7) |
| 8 | CC-Type2-[$L_aI_d$]-L28Y | Ac-G EIAQALK EIAKALK EIAWAL KEIAQAYK G-NH2 | 5.7 | 7nfj (7) |
| 9 | CC-Type2-[$L_aI_d$]-L14A | Ac-G EIAQALK EIAKAAK EIAWALK EIAQALK G-NH2 | 5.8 | 7nfg (6) |
| 10 | CC-Type2-[$L_aI_d$]-I24A | Ac-G EIAQALK EIAKALK EIAWALK EAAQALK G-NH2 | 5.7 | 7nff (8) |
| 11 | CC-Type2-[$L_aI_d$]-I17H | Ac-G EIAQALK EIAKALK EHAWALK EIAQALK G-NH2 | 5.6 | - |
| 12 | CC-Type2-[$L_aI_d$]-L21S-I24Y | Ac-G EIAKALR EIAKALR EIAWASR EYAKALR G-NH2 | 6.8 | - |
| 13 | CC-Type2-[$L_aI_d$]-I24Y | Ac-G EIAQALK EIAKALK EIAWALK EYAQALK G-NH2 | 5.8 | - |
| 14 | CC-Type2-[$L_aI_d$]-L7Y | Ac-G EIAQAYK EIAKALK EIAWALK EIAQALK G-NH2 | 6.0 | 7nfi (8) |
| 15 | CC-Type2-[$L_aI_d$]-L21K | Ac-G EIAKALK EIAKALK EIAWAKK EIAKALK G-NH2 | 7.8 | 7nfm (8) |
| 16 | Trimer | Ac-G EIAAIKQ EIAAIKK EIAAIKW EIAAIKQ G-NH2 | 3.0 | 4dzl (3) |

**Supplementary Table 2. Statistics for the octamer crystal structure.**

| **PDB ID** | **9f5a** |
|---|---|
| **Data Collection** | |
| Source | Diamond I04 |
| Detector | Detector EIGER2 XE 16M |
| Wavelength | 0.975 |
| Resolution range | 34.01 - 2.0 (2.072 - 2.0) |
| Space group | I 4 2 2 |
| Unit cell | 48.0978 48.0978 127.147 90 90 90 |
| Total reflections | 10776 (1022) |
| Unique reflections | 5388 (511) |
| Multiplicity | 2.0 (2.0) |
| Completeness (%) | 99.42 (99.02) |
| Mean I/sigma(I) | 37.47 (8.65) |
| Wilson B-factor | 34.31 |
| R-merge | 0.005393 (0.04893) |
| R-meas | 0.007627 (0.0692) |
| R-pim | 0.005393 (0.04893) |
| CC1/2 | 1 (0.998) |
| CC* | 1 (1) |
| **Refinement** | |
| Reflections used in refinement | 5359 (506) |
| Reflections used for R-free | 538 (51) |
| R-work | 0.2388 (0.2648) |
| R-free | 0.2525 (0.3499) |
| CC(work) | 0.956 (0.957) |
| CC(free) | 0.993 (0.788) |
| Number of non-hydrogen atoms | 482 |
| macromolecules | 441 |
| ligands | 7 |
| solvent | 34 |
| Protein residues | 60 |
| RMS(bonds) | 0.005 |
| RMS(angles) | 0.67 |
| Ramachandran favored (%) | 100 |
| Ramachandran allowed (%) | 0 |
| Ramachandran outliers (%) | 0 |
| Rotamer outliers (%) | 0 |
| Clashscore | 1.1 |
| Average B-factor | 42.18 |
| macromolecules | 41.92 |
| ligands | 50.85 |
| solvent | 43.71 |

**Supplementary Table 3. ISAMBARD modelling scores and parameters for the octamer. *Parameters extracted from the crystal structure.**

| Oligomeric state | Phi-Cα (°) | Radius (Å) | Pitch (Å) | BUDE energy per chain |
|---|---|---|---|---|
| 7 | 113.8 | 9.6 | 308.5 | -560.8 |
| 8 | 113.9 | 10.6 | 377.0 | -583.6 |
| **8* | *121.9* | *11.1* | *556.2* | - |

#### 2.2 TCPCS measurements and fits

**Supplementary Table 4. DPH lifetimes after excitation at 352 nm.** Fluorescence decays were fitted to mono- or bi-exponential decay equations. See Figures S12 and S13.

| | Heptamer | Octamer | Hexamer |
|---|---|---|---|
| $A_1$ | 0.315 | - | - |
| $A_2$ | 0.657 | 0.973 | 0.971 |
| $\tau_1$ | 9.4 ± 0.17 ns | - | - |
| $\tau_2$ | 18.6 ± 0.17 ns | 15.8 ± 0.17 ns | 19.8 ± 0.17 ns |
| $\tau_{average}$ | **15.9 ± 0.17 ns** | **15.8 ± 0.17 ns** | **19.8 ± 0.17 ns** |

**Supplementary Table 5. Nile Red lifetimes after excitation at 535 nm.** Fluorescence decays were fitted to mono- or bi-exponential decay equations. See Figures S14 and S15.

| | Heptamer | Octamer | Hexamer |
|---|---|---|---|
| $A_1$ | - | 0.264 | - |
| $A_2$ | 1.039 | 0.758 | 1.086 |
| $\tau_1$ | - | 1.9 ± 0.17 ns | - |
| $\tau_2$ | 4.1 ± 0.17 ns | 5.0 ± 0.17 ns | 4.9 ± 0.17 ns |
| $\tau_{average}$ | **4.1 ± 0.17 ns** | **4.2 ± 0.17 ns** | **4.9 ± 0.17 ns** |

**Supplementary Table 6. Nile Red lifetimes after excitation at 535 nm with DPH.** Fluorescence decays were fitted to mono- or bi-exponential decay equations. See Figures S14 and S16.

| | Heptamer | Octamer | Hexamer |
|---|---|---|---|
| $A_1$ | - | - | 0.372 |
| $A_2$ | 1.039 | 1.042 | 0.747 |
| $\tau_1$ | - | - | 1.0 ± 0.17 ns |
| $\tau_2$ | 4.9 ± 0.17 ns | 5.1 ± 0.17 ns | 4.9 ± 0.17 ns |
| $\tau_{average}$ | **4.9 ± 0.17 ns** | **5.1 ± 0.17 ns** | **4.0 ± 0.17 ns** |

**Supplementary Table 7. Simultaneous FRET lifetime fits from DPH and Nile Red channels after excitation at 352 nm.** Floated parameters in bold. Fluorescence decays were fitted to bi-exponential decay equations while keeping average DPH and Nile Red lifetimes from Tables S4 and S6 fixed. See Figure S17.

| | Heptamer | Octamer |
|---|---|---|
| $\boldsymbol{A_1}$**(DPH)** | **0.086** | **0.149** |
| $\boldsymbol{A_2}$**(DPH)** | **0.898** | **0.837** |
| $\boldsymbol{A_1}$(NR) | **–0.310** | **–0.337** |
| $\boldsymbol{A_2}$**(NR)** | **1.284** | **1.289** |
| $\boldsymbol{\tau_{FRET}}$ | **0.98 ± 0.17 ns** | **1.3 ± 0.17 ns** |
| $\tau_{DPH}$ | 15.9 ± 0.17 ns | 15.8 ± 0.17 ns |
| $\tau_{NR}$ | 4.9 ± 0.17 ns | 5.1 ± 0.17 ns |

#### 2.3 TA measurements and fits

**Supplementary Table 8. FRET lifetime fits from Nile Red channel after excitation at 352 nm.** Floated parameters in bold. Average Nile Red decay lifetime from Table S6 was kept fixed. See Figure S22.

| | Heptamer | Octamer |
|---|---|---|
| $\boldsymbol{A_1}$ | **0.0015** | **0.00012** |
| $\boldsymbol{A_2}$ | **-0.0025** | **-0.00020** |
| $\boldsymbol{\tau_1}$ | **10.4 ± 2.4 ps** | **7.9 ± 5.7 ps** |
| $\tau_2$ | 4.9 ± 0.17 ns | 5.1 ± 0.17 ns |

#### 2.4 FRET distance calculation

**Supplementary Table 9. DPH fluorescence quantum yield ($\Phi_F$) measurements** in absence of Nile Red **for the peptides in this study and in buffer.**

| | $\Phi_F$ |
|---|---|
| Hexamer | 0.63 ± 0.01 |
| Heptamer | 0.65 ± 0.02 |
| Octamer | 0.39 ± 0.02 |
| Buffer | 0.08 ± 0.03 |

**Supplementary Table 10. Förster spectral overlap between DPH and Nile Red.**

| Heptamer | Octamer |
|---|---|
| $8.5 \times 10^{14}$ $nm^4$/(M cm) | $9.5 \times 10^{14}$ $nm^4$/(M cm) |

#### 3 Supplementary Figures

##### 3.1 Biophysical characterisation

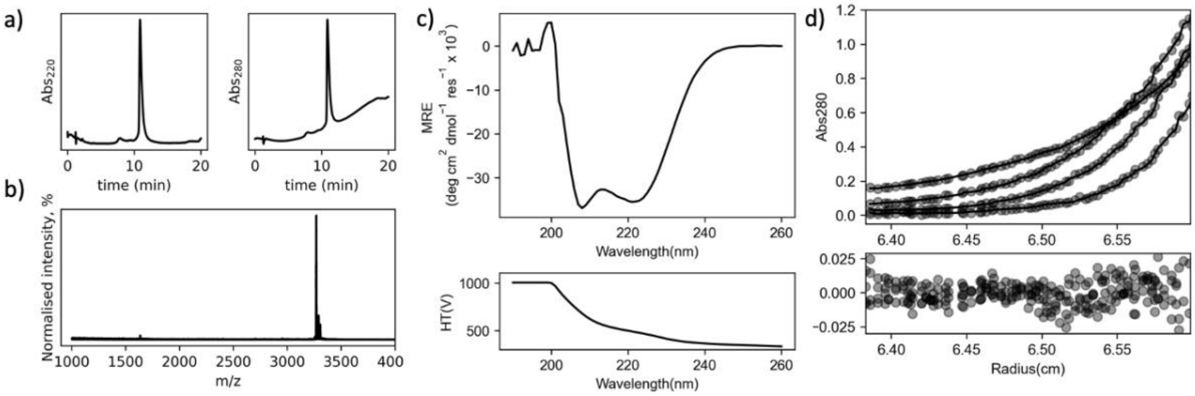

**Supplementary Figure 1. Biophysical characterisation of the octamer**. **a,** Analytical HPLC and **b,** MALDI-TOF traces. **c,** CD spectra. Conditions: 100 μM peptide, HEPES, pH 7, 20 °C. **d,** Sedimentation equilibrium traces. Fitted MW = 23.9 ± 0.3 kDa (7.4 x monomer, 95% confidence limits). Data were collected from 24 to 48 krpm at 6 krpm intervals. Fitted single-ideal species model curves are overlaid in solid line. Conditions: 75 μM peptide, HEPES, pH 7, 20 °C. Although sedimentation equilibrium returned 7.4 x monomer, parametric modelling predicted an octameric assembly (Table S3). We also observe an octameric assembly in a crystal state (Table S2, PDB: 9a5f).

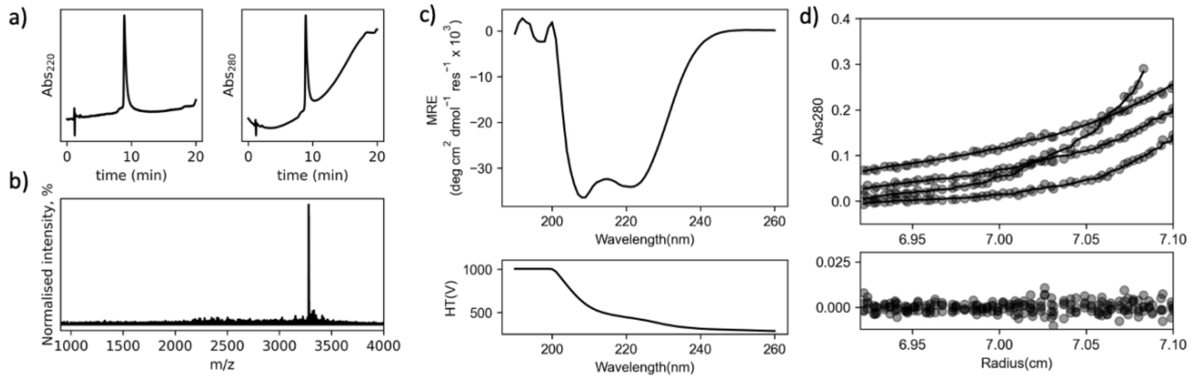

**Supplementary Figure 2. Biophysical characterisation of CC-Type2-[$I_aI_d$]- I17V-I21Y**. **a,** Analytical HPLC and **b,** MALDI-TOF traces. **c,** CD spectra. Conditions: 100 μM peptide, HEPES, pH 7, 20 °C. **d,** Sedimentation equilibrium traces. Fitted MW = 23.9 ± 0.3 kDa (7.0 x monomer, 95% confidence limits). Data were collected from 25 to 45 krpm at 5 krpm intervals. Fitted single-ideal species model curves are overlaid in solid line. Conditions: 75 μM peptide, HEPES, pH 7, 20 °C.

#### 3.2 DPH FRET acceptor screen

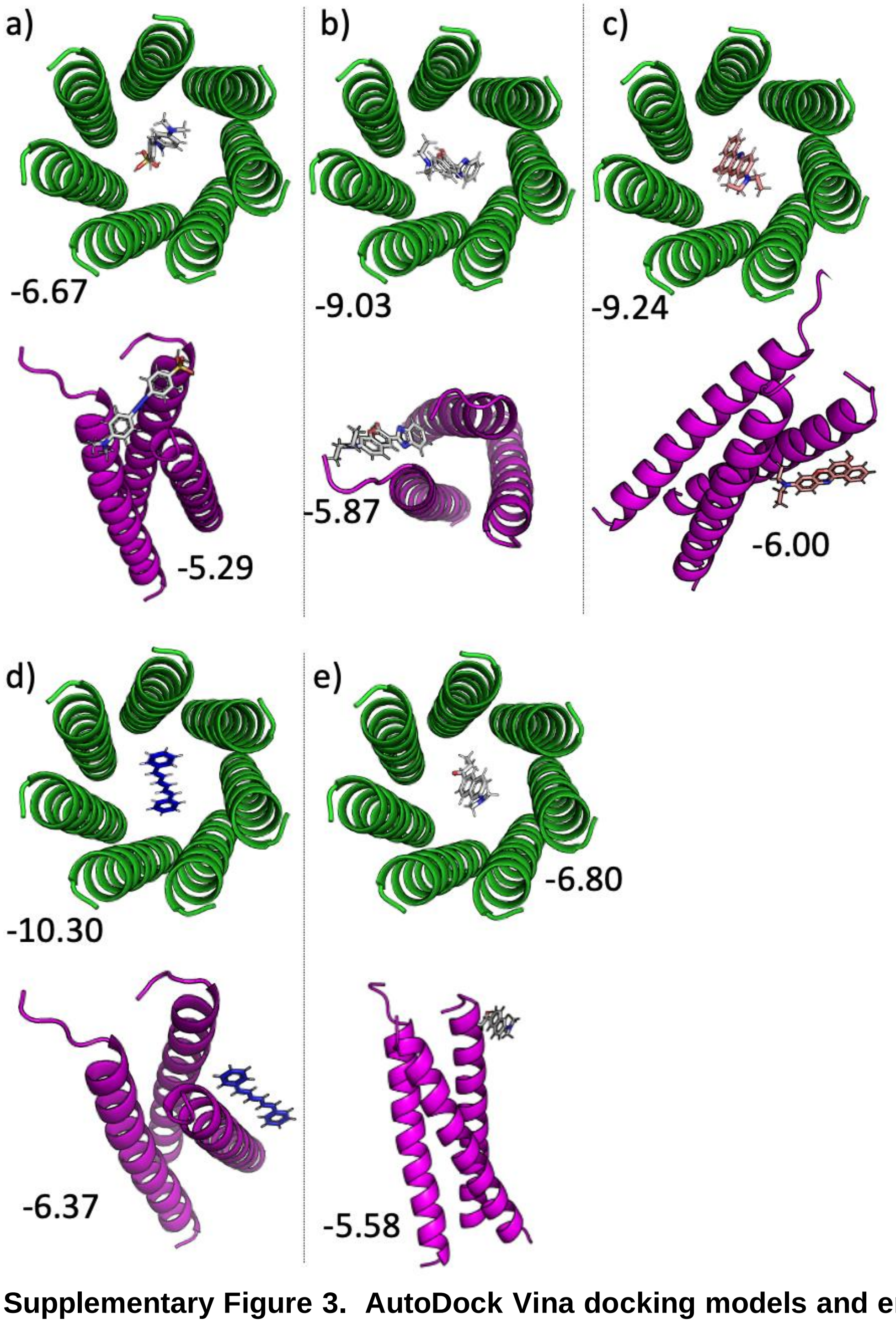

**Supplementary Figure 3. AutoDock Vina docking models and energies (kcal/mol) of potential FRET acceptors (a) Methyl Orange, (b) Coumarin-7 and (c) Nile Red compared to know binders (d) DPH and (e) Prodan.** Receptors: the heptamer (PDB: 6g66, green) and a negative control CC-Tri (PDB: 4dzl, magenta). Ligands were geometry-optimised with a MOPAC (PM6-D3H4, COSMO) and docked with AutoDock Vina 1.2 with exhaustiveness of 64. DPH torsion angles were set to 0.

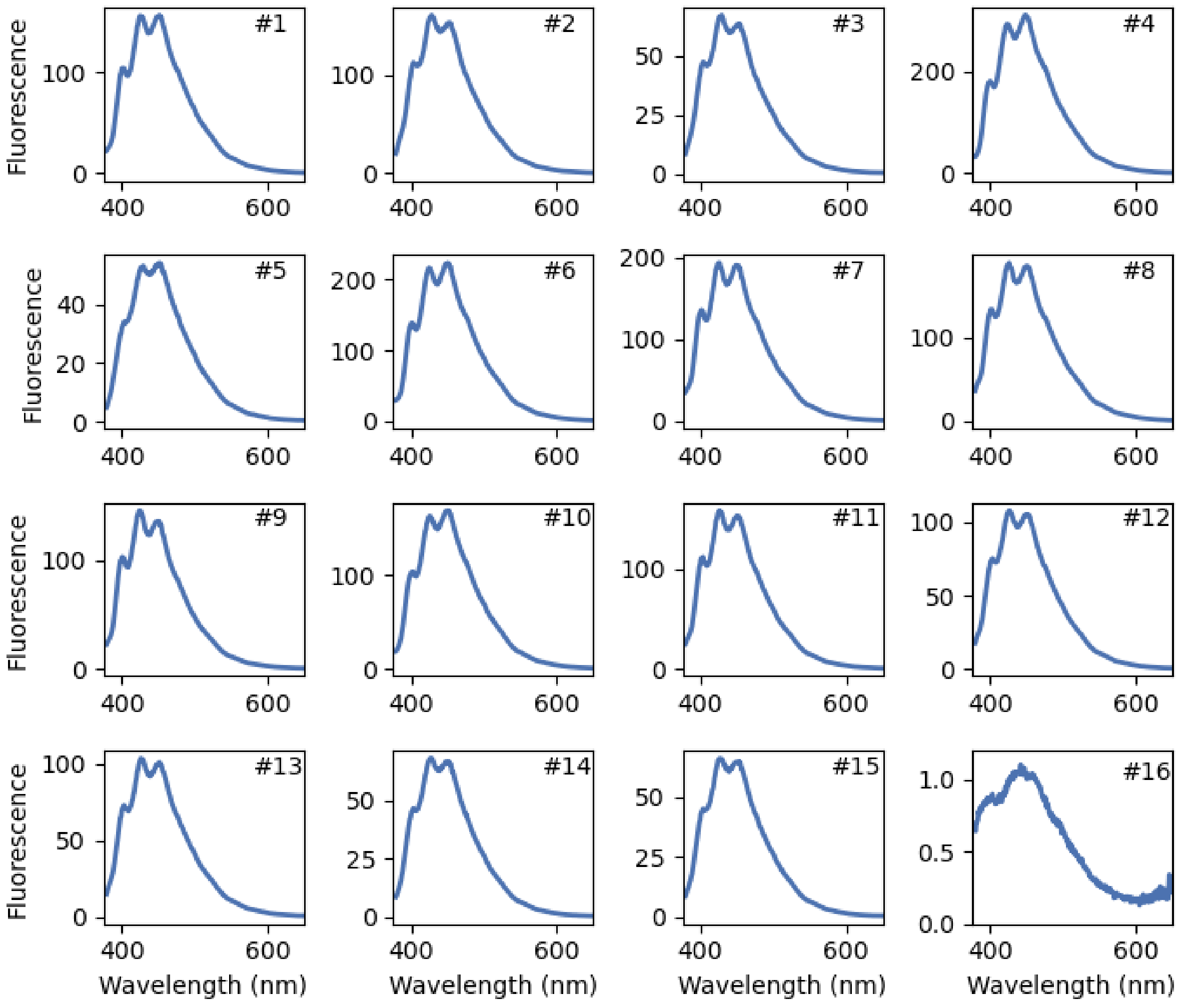

**Supplementary Figure 4. Fluorescence spectra of DPH binding to αHBs with hexameric to octameric oligomeric states and lumenal hydrophobic and/or aromatic residues.** #1 is the heptamer, #2 is the octamer and #4 is the hexamer. Normalised to the trimer control (#16). See Table S1 for sequences and biophysical characterisation. Conditions: 3 μM DPH, 5 μM peptide assembly, HEPES, 10% v/v MeCN, pH 7, 352 nm excitation.

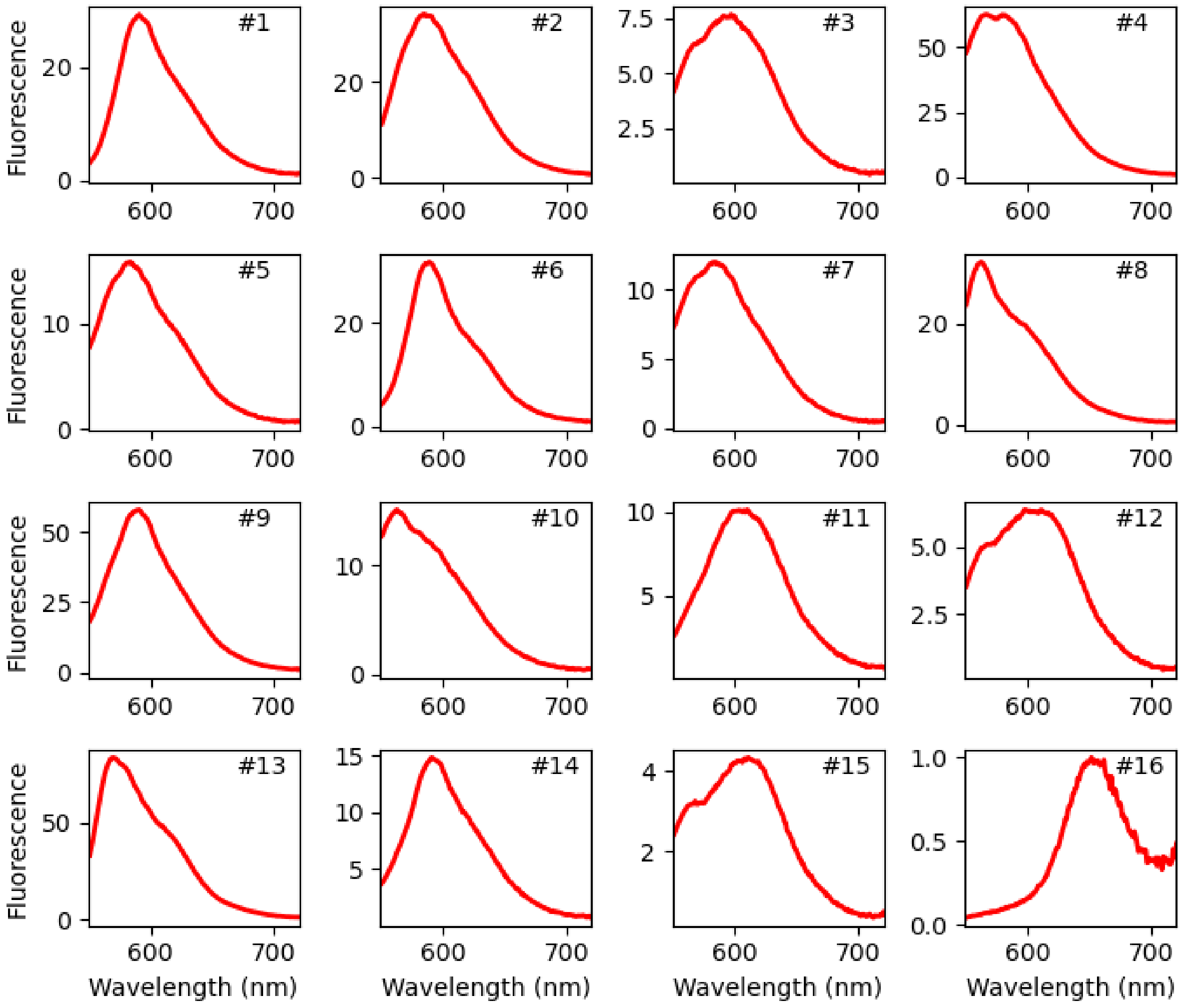

**Supplementary Figure 5. Fluorescence spectra of Nile Red binding to αHBs with hexameric to octameric oligomeric states and lumenal hydrophobic and/or aromatic residues.** #1 is the heptamer, #2 is the octamer and #4 is the hexamer. Normalised to the trimer control (#16). See Table S1 for sequences and biophysical characterisation. Conditions: 3 μM Nile Red, 5 μM peptide assembly, HEPES, 10% v/v MeCN, pH 7, 520 nm excitation.

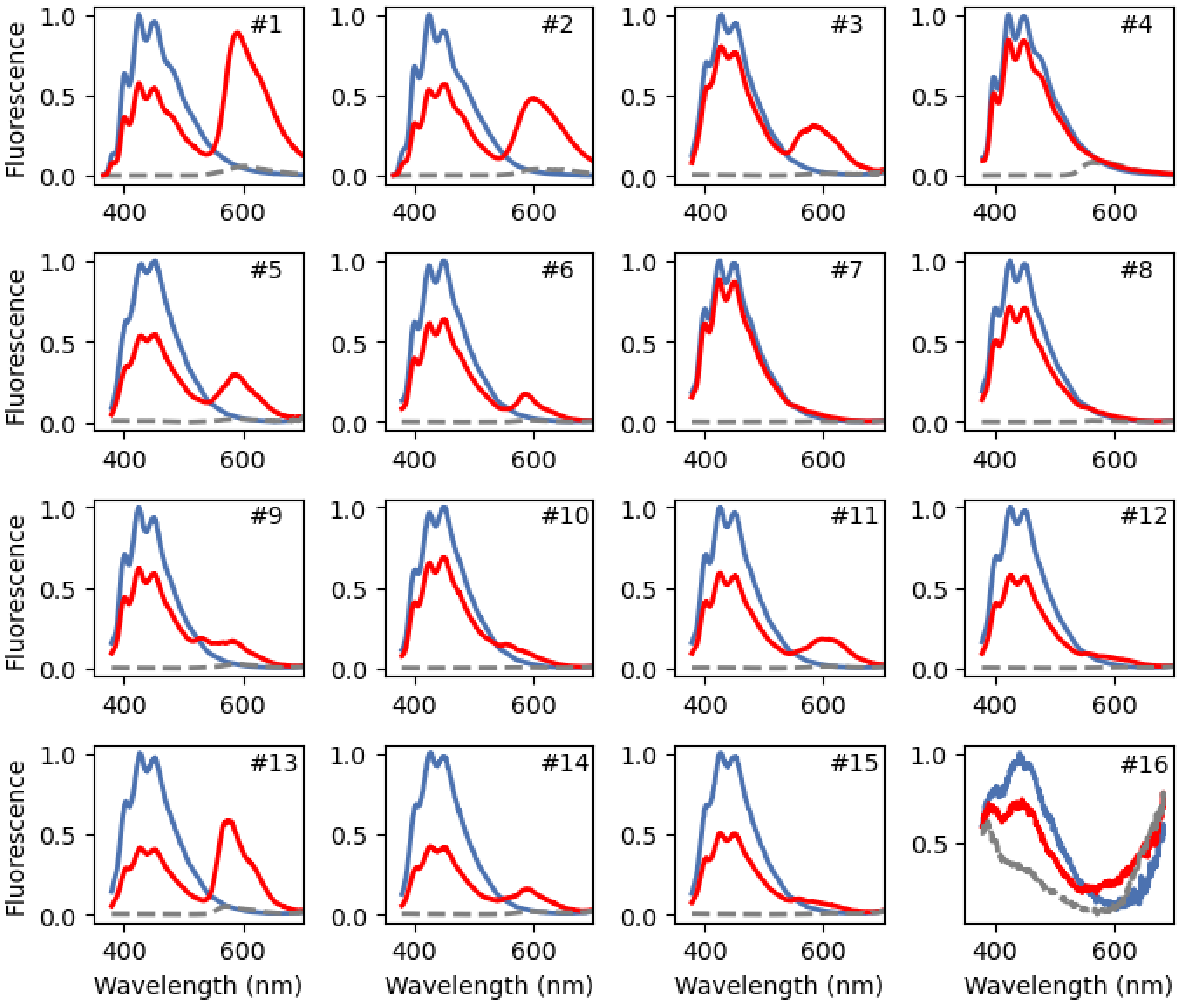

**Supplementary Figure 6. Steady-state Fluorescence Spectra reveals FRET between DPH and Nile Red within aHBs with hexameric to octameric oligomeric states and lumenal hydrophobic and/or aromatic residues.** Blue – DPH only, dashed grey – Nile Red only, red – DPH and Nile Red. Normalised to DPH emission without Nile Red. #1 is the heptamer, #2 is the octamer and #4 is the hexamer. #16 is the trimer control. Conditions: 3 μM Nile Red (red and grey), 3 μM DPH (blue and red), 5 μM peptide assembly, HEPES, 10% v/v MeCN, pH 7, 352 nm excitation. Table S1 for sequences and biophysical characterisation.

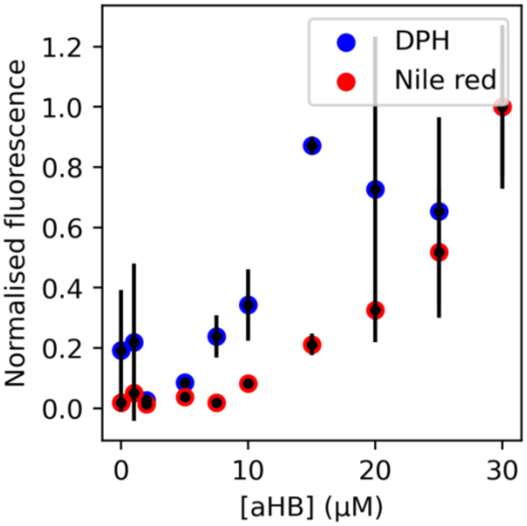

**Supplementary Figure 7. Titration curves showing non-specific DPH and Nile Red interactions with the trimer.** Conditions: 0.5 μM DPH or Nile Red, 0 – 30 μM peptide assembly, HEPES, 10% v/v MeCN, pH 7. The data are the mean of three independent repeats, error bars represent the standard deviation from the mean.

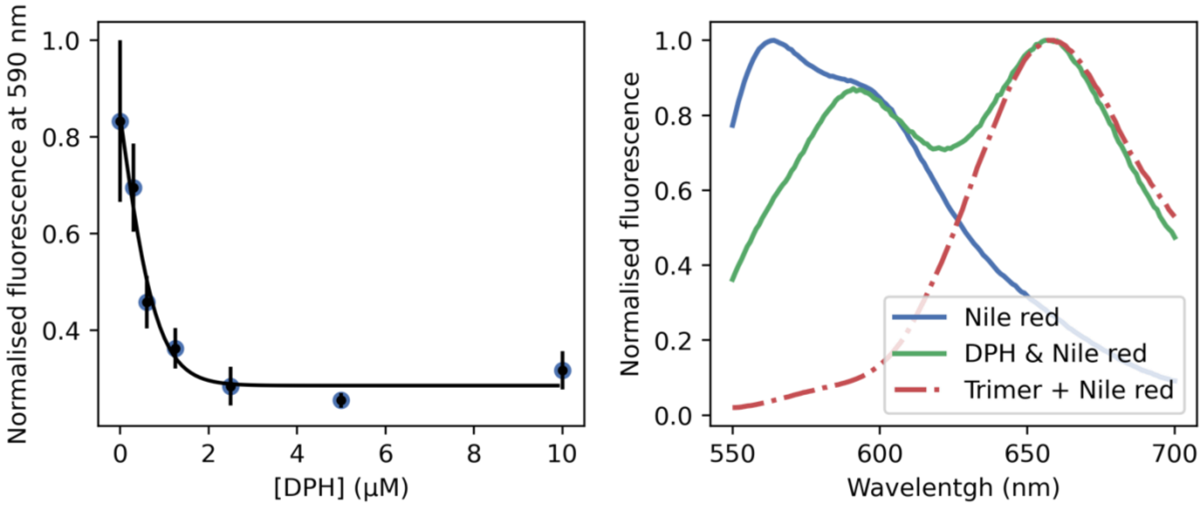

**Supplementary Figure 8. Nile Red displacement by DPH in the hexamer. a,** Titration curve (blue) and one-site displacement model fit (black, $K_i$ = 0.3 ± 0.4 μM). Conditions: 0.5 μM Nile Red, 5 μM peptide assembly, DPH varied from 0 to 5 μM, HEPES, 10% v/v MeCN, pH 7. **b,** Nile Red spectra sift upon DPH addition. The trimer is a closed bundle which shows only non-specific Nile Red binding. Conditions: 3 μM Nile Red, 3 μM DPH (green and red), 5 μM peptide assembly, HEPES, 10% v/v MeCN, pH 7, 520 nm excitation. The data are the mean of three independent repeats, error bars represent the standard deviation from the mean.

##### 3.3 Calculation of FRET distance

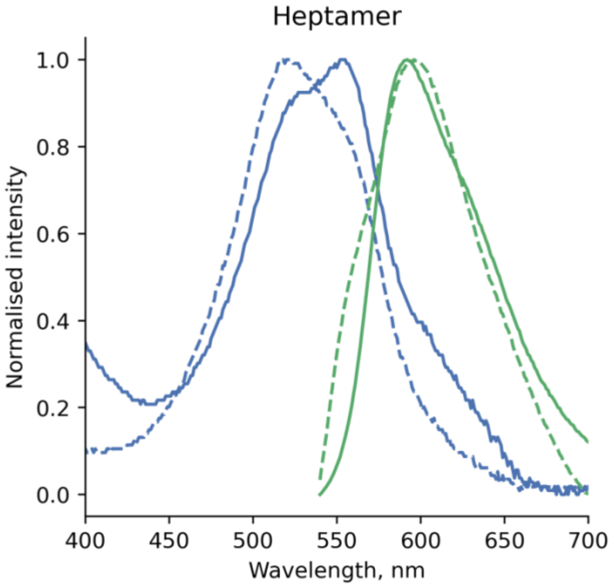

**Supplementary Figure 9. Nile Red absorption (blue) and emission (green, $\lambda_{ex}$ = 520 nm) spectra with 3 µM DPH (solid) and without (dashed).** Conditions: 3 µM Nile Red, 3 µM DPH (solid only), 5 µM peptide assembly, HEPES, 10% v/v MeCN, pH 7.

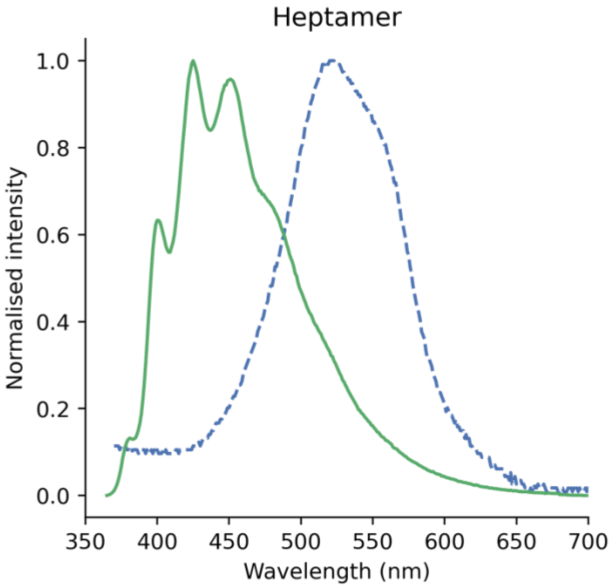

**Supplementary Figure 10. Normalised Nile Red absorption (blue) and DPH emission (green) spectra used for calculating the spectral overlap for peptides in this study.** Conditions: 3 µM Nile Red or 3 µM DPH, 5 µM peptide assembly, HEPES, 10% v/v MeCN, pH 7, 352 nm excitation (DPH emission).

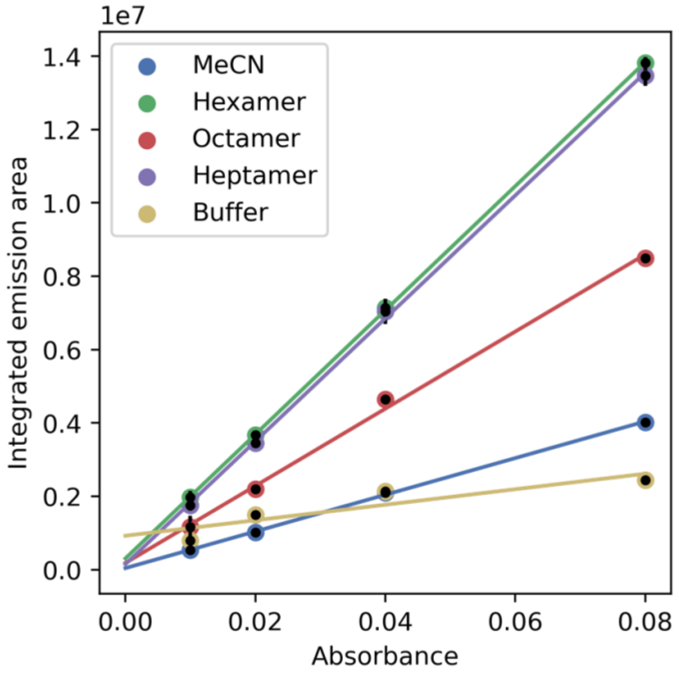

**Supplementary Figure 11. Relative DPH quantum yield (Φ) measurements for the peptides in this study.** Reference: DPH in acetonitrile (Φ = 0.19). Also see Table S9. Conditions: 0.01 – 0.08 µM DPH, 5 µM peptide assembly, HEPES, 10% v/v MeCN, pH 7, 352 nm excitation. The data are the mean of three independent repeats, error bars represent the standard deviation from the mean.

#### 3.4 TCSPC measurements and fits

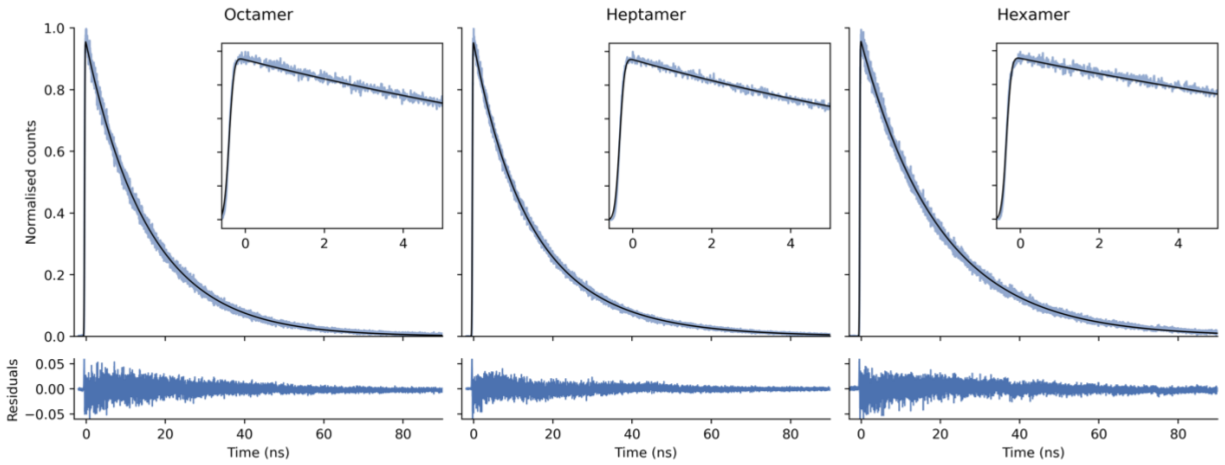

**Supplementary Figure 12. TCSPC traces and fits showing DPH fluorescence decay for peptides in this study.** Zoomed in view shows the first 5 ns. Fit parameters are shown in Table S4, time-resolved emission spectra and the data range used for fitting shown in Figure S10. Conditions: 3 µM DPH, 5 µM peptide assembly, HEPES, 10% v/v MeCN, pH 7, 352 nm excitation. Time-resolved emission spectra was acquired with the GEMINI interferometer inline.

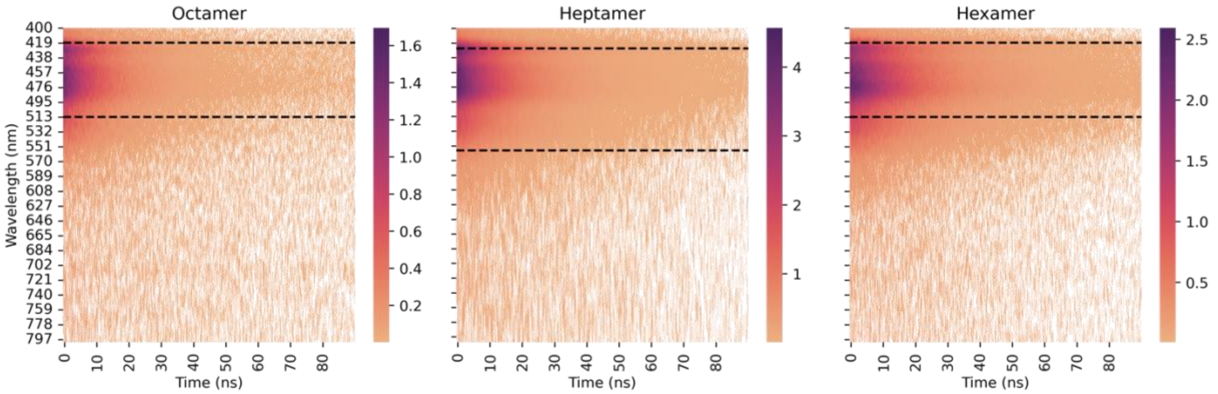

**Supplementary Figure 13. Time-resolved emission spectra derived from the interferometric map showing DPH fluorescence decay for peptides in this study.** Data range used for traces in Figure S12 shown in dashed lines. Conditions: 3 µM DPH 5 µM peptide assembly, HEPES, 10% v/v MeCN, pH 7, 352 nm excitation.

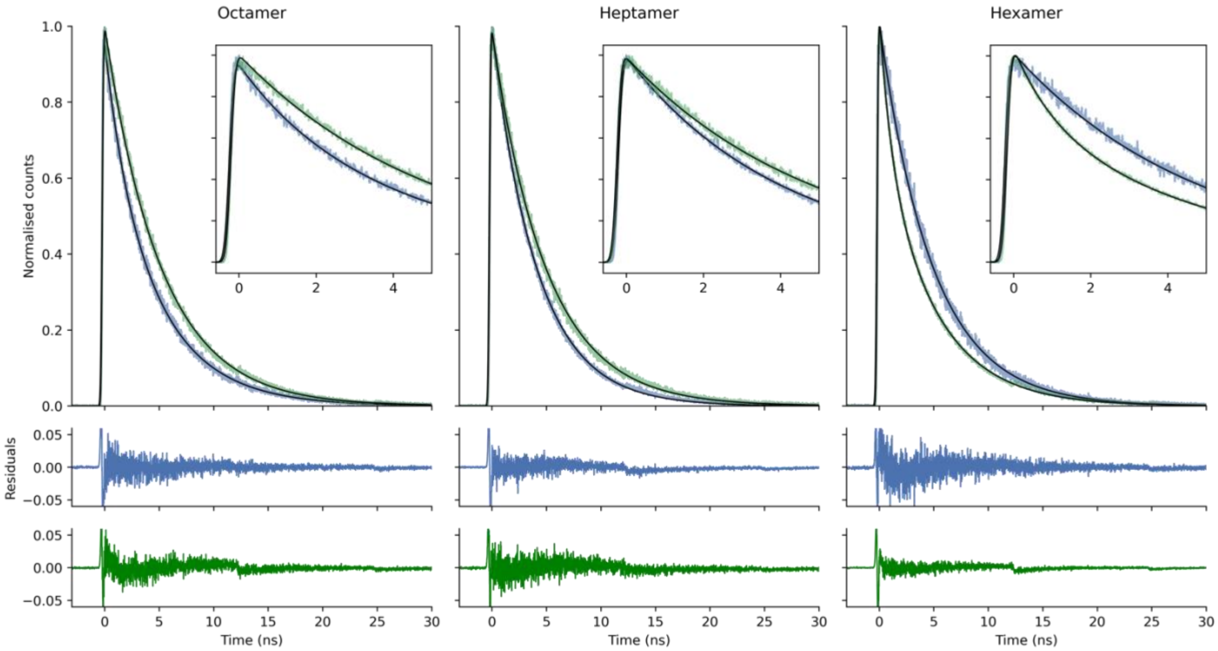

**Supplementary Figure 14. TCSPC traces and fits showing Nile Red fluorescence decay with DPH (green) or without DPH (blue) for peptides in this study.** Zoomed in view shows the first 5 ns. Fit parameters are shown in Tables S5 - S6, time-resolved emission spectra and the data range used for fitting shown in Figures S15 - S16. Conditions: 3 µM Nile Red, 3 µM DPH (green only), 5 µM peptide assembly, HEPES, 10% v/v MeCN, pH 7, 535 nm excitation. For the hexamer with DPH and Nile Red, TCSPS spectra was collected with a >565 nm filter due to a weaker emission. For the other samples, time-resolved emission spectra were acquired the GEMINI interferometer inline.

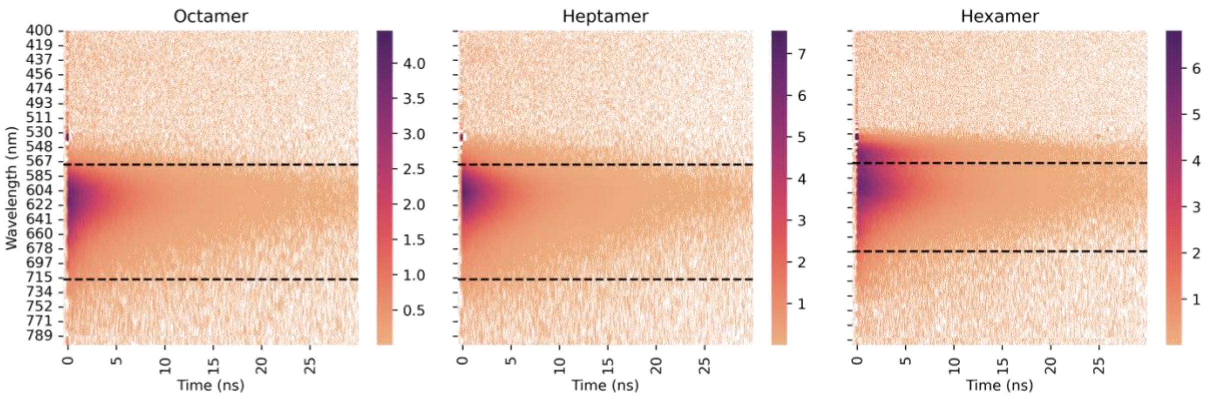

**Supplementary Figure 15. Time-resolved emission spectra derived from the interferometric map showing Nile Red fluorescence decay without DPH for peptides in this study.** Data range used for traces in Figure S14 shown in dashed lines. Conditions: 3 μM Nile Red, 5 μM peptide assembly, HEPES, 10% v/v MeCN, pH 7, 535 nm excitation.

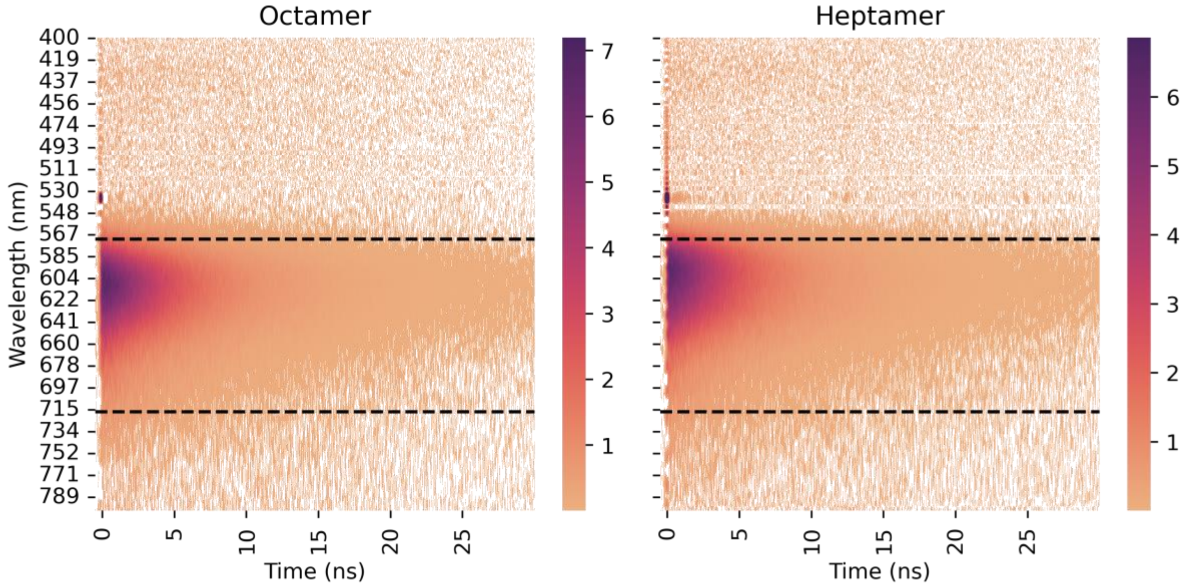

**Supplementary Figure 16. Time-resolved emission spectra derived from the interferometric map showing Nile Red fluorescence decay with DPH for peptides in this study.** Data range used for traces in Figure S14 shown in dashed lines. Time-resolved emission spectra for the hexamer with DPH and Nile Red were not collected due to a weaker signal (Nile Red displacement). Conditions: 3 μM Nile Red, 3 μM DPH, 5 μM peptide assembly, HEPES, 10% v/v MeCN, pH 7, 535 nm excitation.

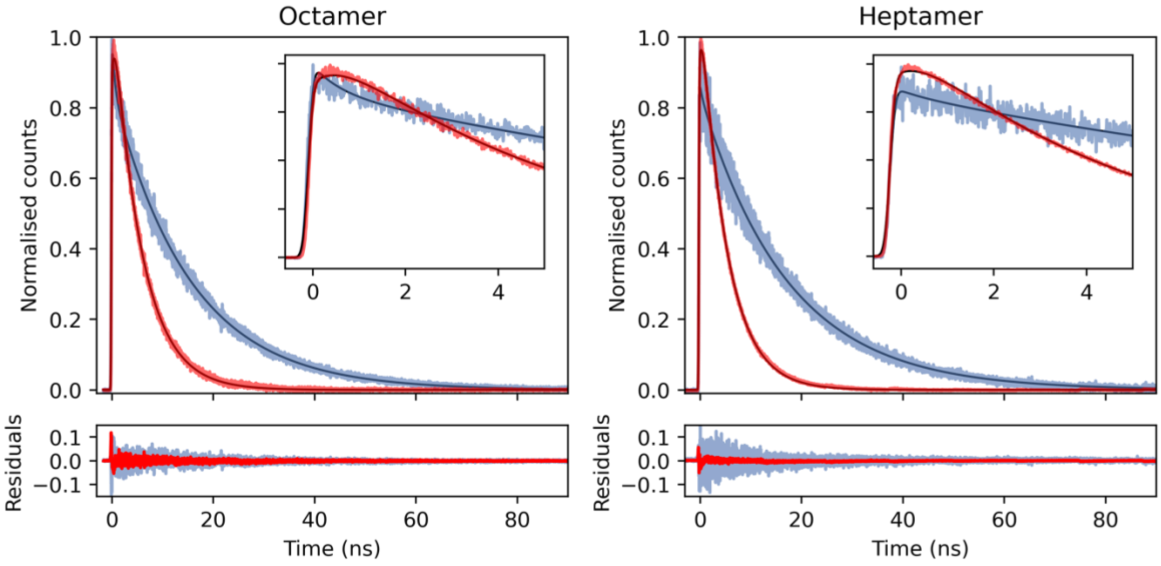

**Supplementary Figure 17. TCSPC traces and fits showing Nile Red (red) and DPH (blue) energy transfer and fluorescence decay for peptides in this study.** The resulting data were fitted simultaneously with a biexponential function convolved with the instrument response for both FRET and intrinsic fluorescence channels while fixing the fluorescence decay constants for DPH and Nile Red to those determined in control measurements. For fitted parameters, see Table S7. Time-resolved emission spectra and the data range used for fitting shown in Figure S18. Conditions: 3 μM Nile Red, 3 μM DPH, 5 μM peptide assembly, HEPES, 10% v/v MeCN, pH 7, 352 nm excitation.

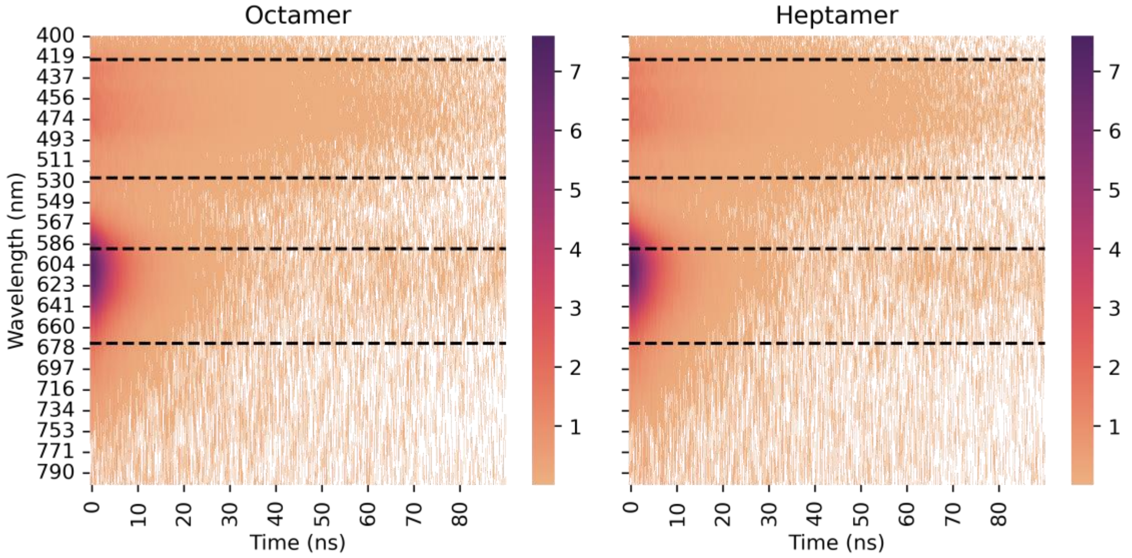

**Supplementary Figure 18. Time-resolved emission spectra derived from the interferometric map showing DPH and Nile Red fluorescence decay for peptides in this study.** Data range used for traces in Figure S17 shown in dashed lines. Conditions: 3 μM Nile Red, 3 μM DPH, 5 μM peptide assembly, HEPES, 10% v/v MeCN, pH 7, 352 nm excitation.

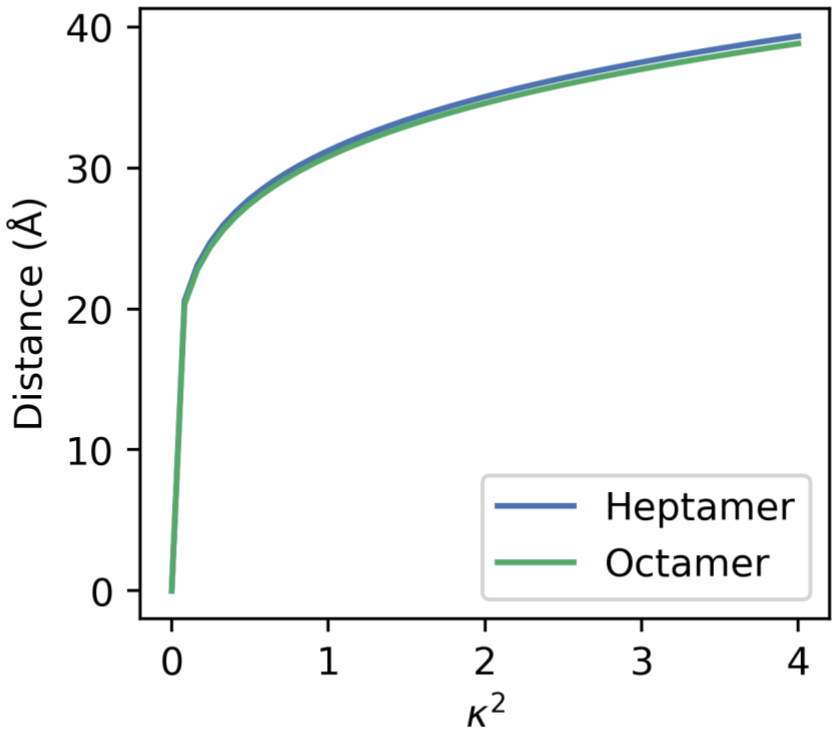

**Supplementary Figure 19. Distance between DPH and Nile red dependence on the orientation factor $\kappa^2$.** The distance was calculated using equation S3 from the FRET lifetimes, see Figure S18 and Table S7.

##### 3.5 TA measurements and fits

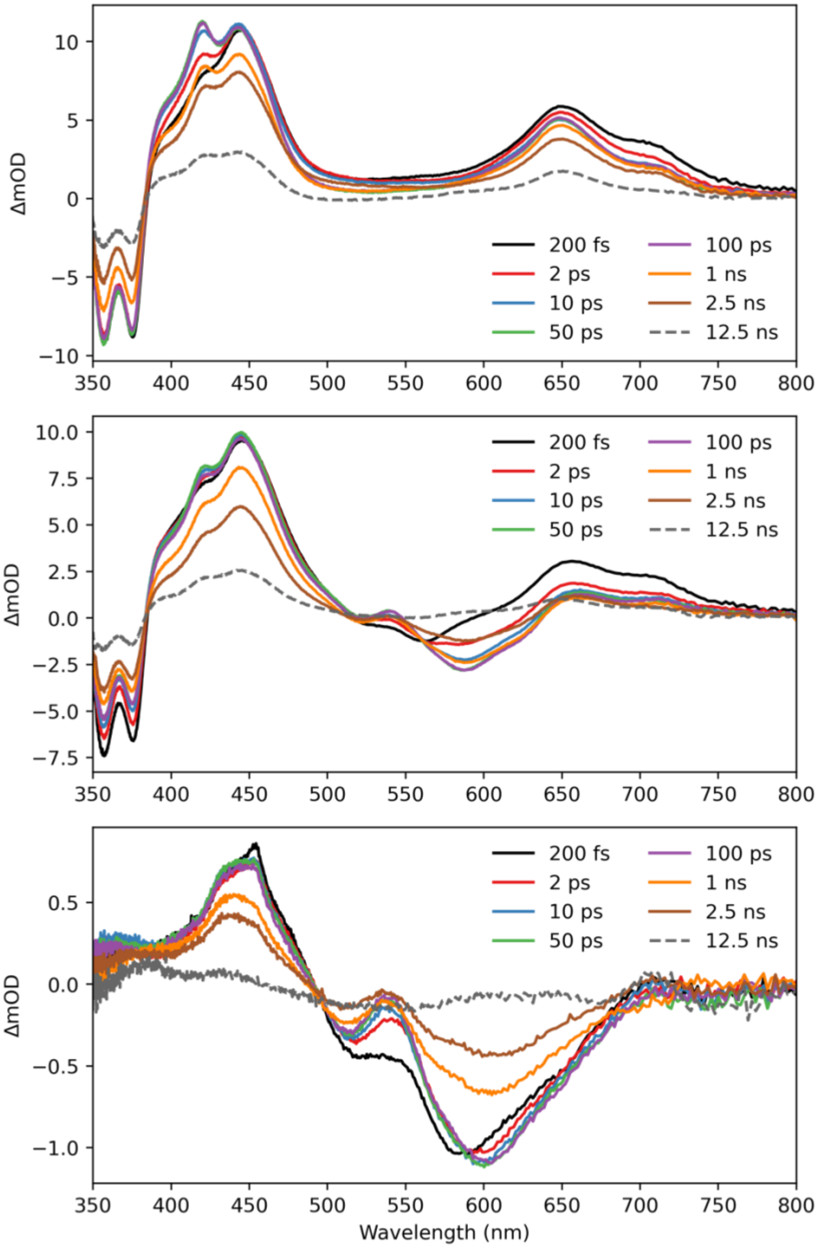

**Supplementary Figure 20. Wavelength resolved transient absorption of the heptamer for DPH and Nile Red for 8 different pump-probe time delays. a,** DPH after 352 nm excitation. **b,** DPH and Nile Red after 352 nm excitation. **c,** Nile Red after 535 nm excitation. TA data slices in (a) and (b) were dominated by positive excited state absorption signals associated with the generation of photoexcited DPH (400 – 500 nm and 650 – 770 nm). Negative signals evident between 350 – 380 nm corresponded to the DPH ground state bleach. The negative signal centered at 590 nm is associated with stimulated emission from Nile Red, and in (b) arises from FRET, and (c) from directed excitation of Nile Red. Conditions: 10 μM DPH and Nile Red, 15 μM peptide assembly, HEPES, 10% v/v MeCN, pH 7. Conditions: 10 μM Nile Red, 10 μM DPH, 15 μM peptide assembly, HEPES, 10% v/v MeCN, pH 7.

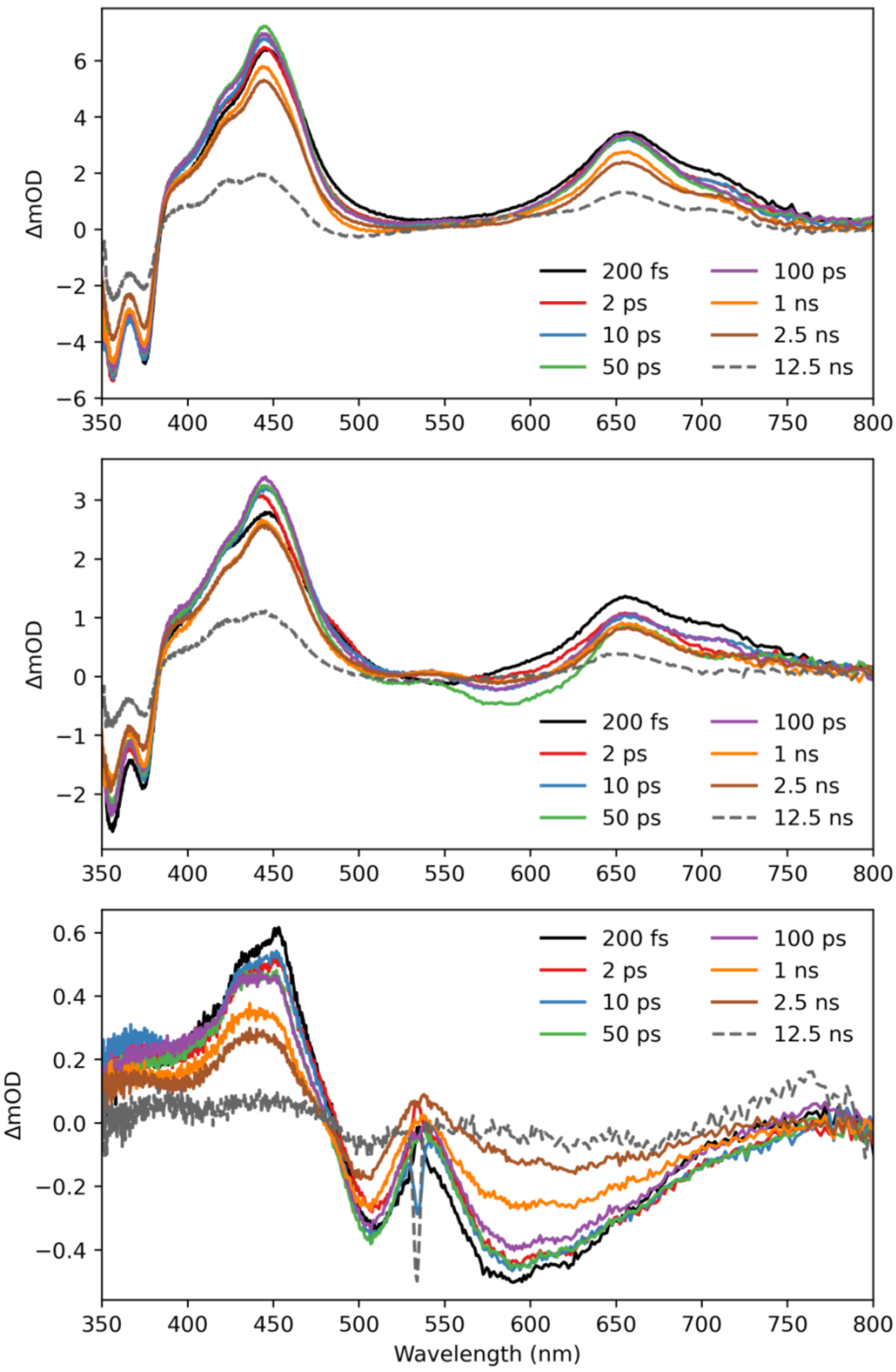

**Supplementary Figure 21. Wavelength resolved transient absorption of the octamer for 8 different pump-probe time delays. a,** DPH after 352 nm excitation. **b,** DPH and Nile Red after 352 nm excitation. **c,** Nile Red after 535 nm excitation. The spectral assignemnts for the octamer data were the same as for the heptamer (see Figure S20). Spike at 535 nm in panel (c) arises from a small amount of pump scatter contamination. Conditions: 10 μM DPH and Nile Red, 15 μM peptide assembly, HEPES, 10% v/v MeCN, pH 7. Conditions: 10 μM Nile Red, 10 μM DPH, 15 μM peptide assembly, HEPES, 10% v/v MeCN, pH 7.

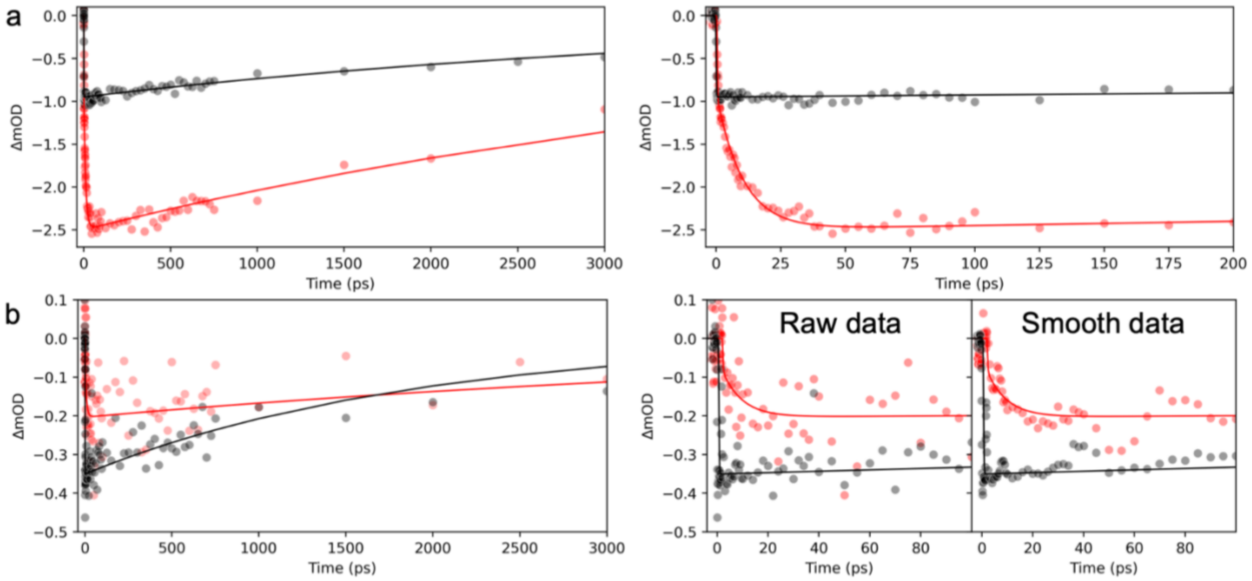

**Supplementary Figure 22. Kinetics obtained by integration over the Nile Red stimulated emission signal for (a) the heptamer and (b) the octamer with Nile Red (535 nm excitation, black circles) and DPH and Nile Red (352 nm excitation, red circles).** Early time dynamics are shown on the right. Fits are shown as solid lines. For fitted parameters, see Table S8. For the octamer we observe a weaker signal due to photodegradation of the peptide. Hence, data smoothed with a moving average of 3 data points is provided to better visualise the exponential growth on the Nile Red stimulated emission. Conditions: 10 μM Nile Red, 10 μM DPH, 15 μM peptide assembly, HEPES, 10% v/v MeCN, pH 7.

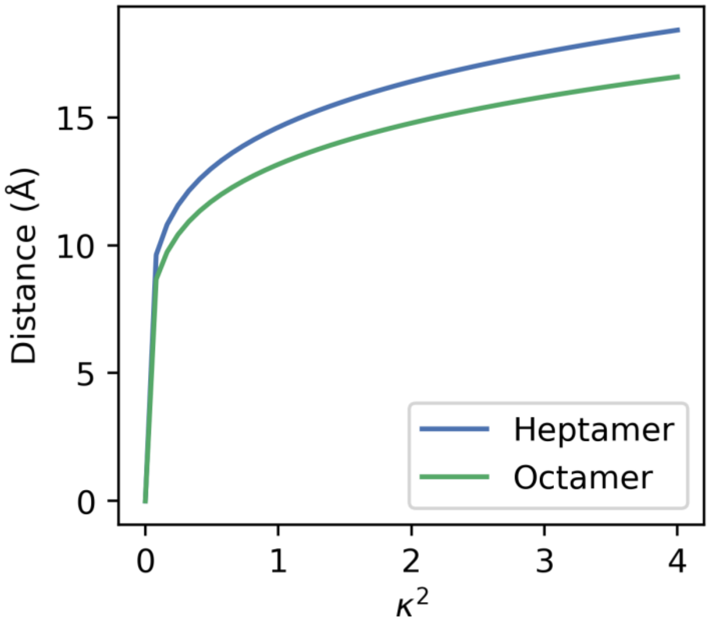

**Supplementary Figure 23. Distance between DPH and Nile red dependence on the orientation factor $\kappa^2$.** The distance was calculated using equation S3 from the FRET lifetimes, see Figure S24 and Table S8.

##### 3.6 Docking and MD simulations

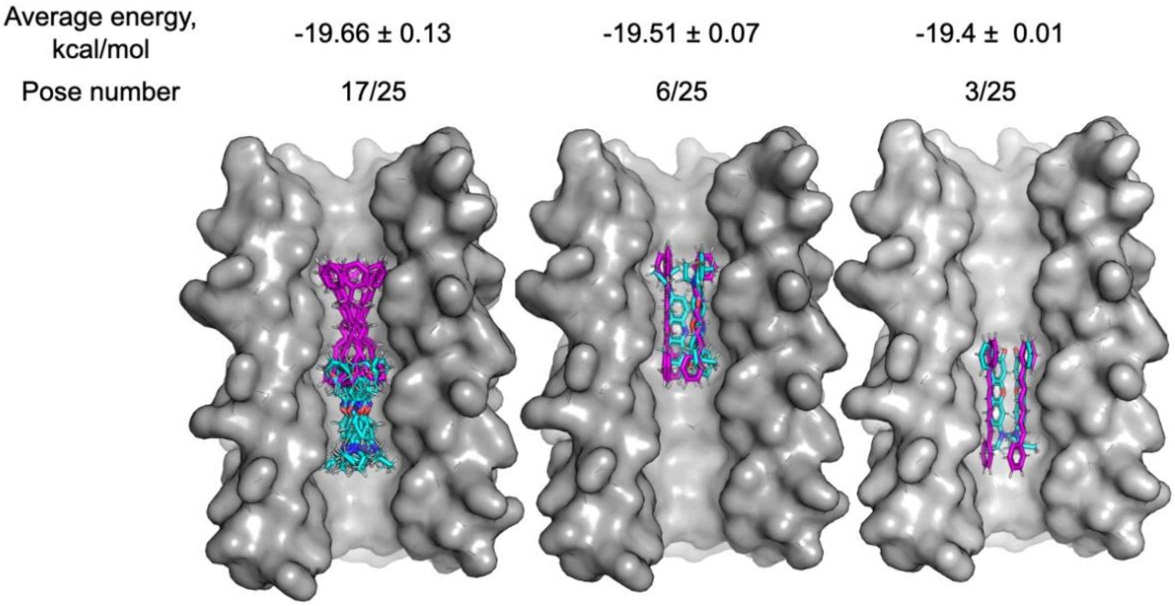

**Supplementary Figure 24. Top 25 AutoDock Vina binding poses for DPH (purple) and Nile Red (cyan) in the heptamer (PDB: 6g66).** Ligands were geometry-optimised with a MOPAC (PM6-D3H4, COSMO) and simultaneously docked with AutoDock Vina 1.2 with exhaustiveness of 64. DPH torsion angles were set to 0.

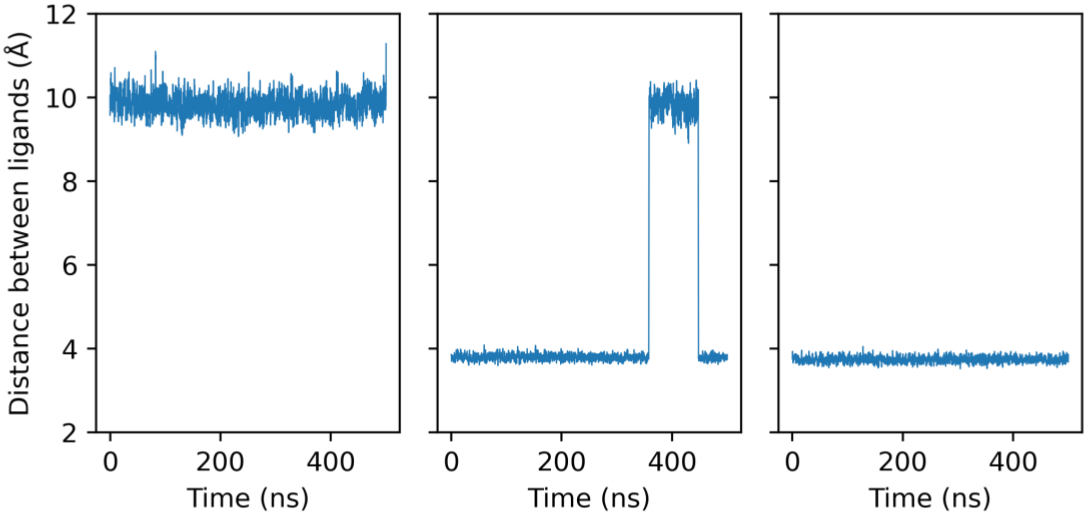

**Supplementary Figure 25. Distance between DPH and Nile red within the heptamer.** 3 x 500ns runs from starting poses in Figure S24. The distance was calculated between the centres of mass of the molecules.

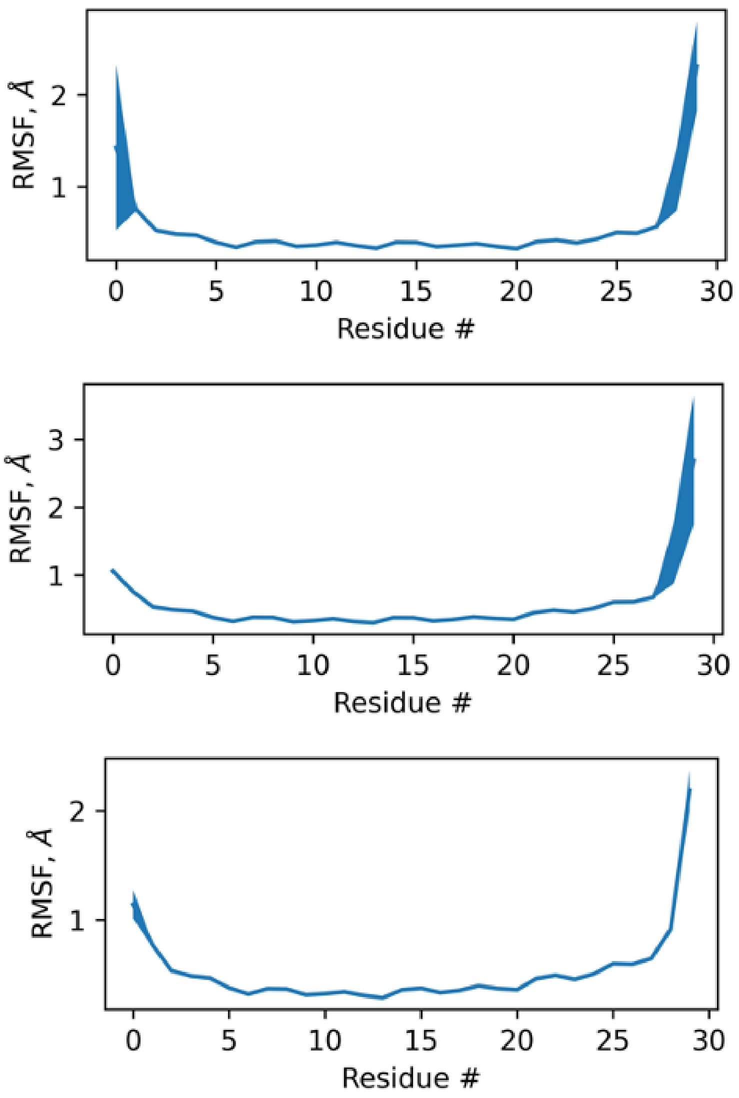

**Supplementary Figure 26. Cα root mean square fluctuation in the heptamer.** Line area shows mean across all chains ± one standard deviation. 3 x 500ns runs from starting poses in Figure S24.

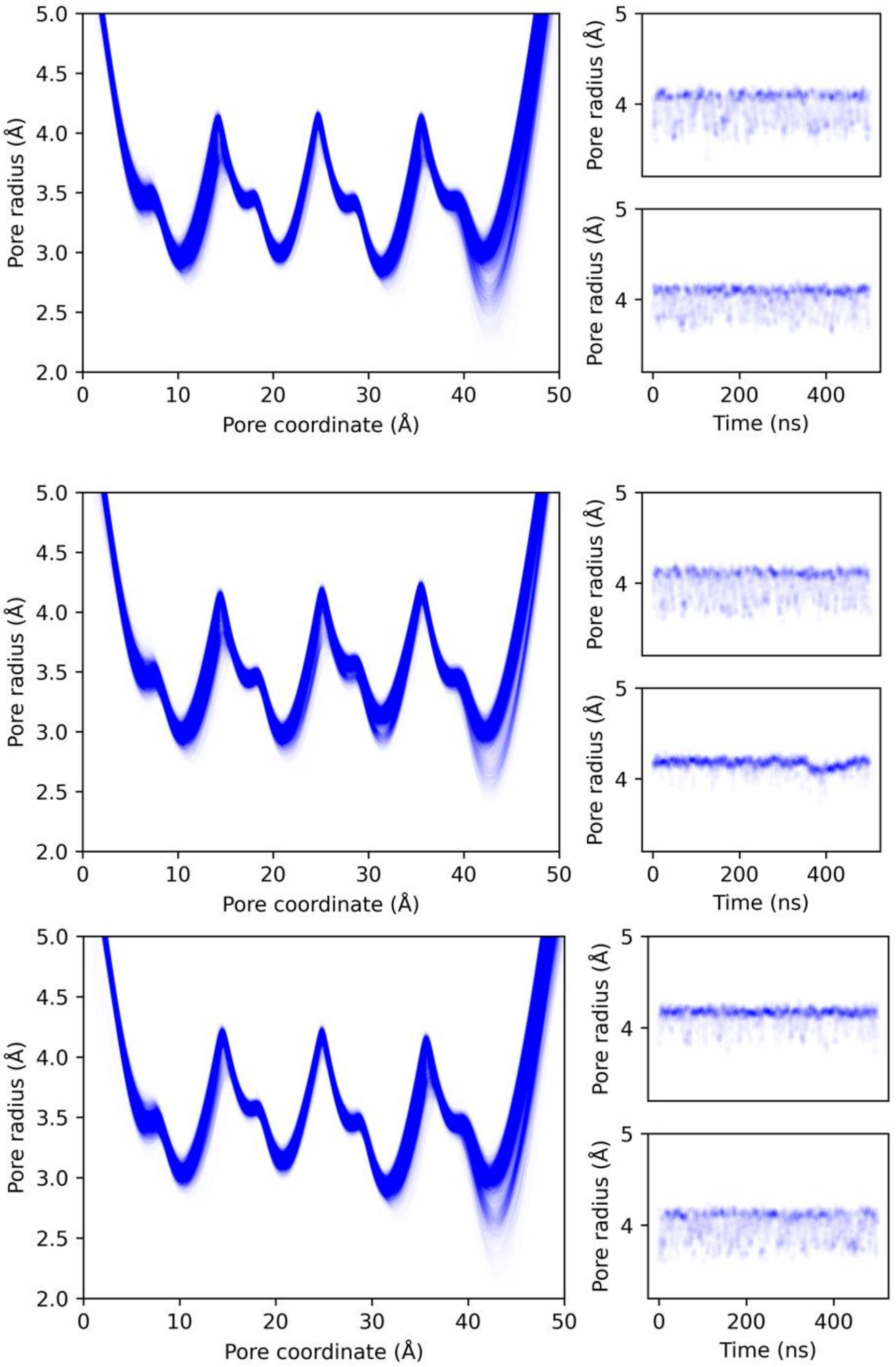

**Supplementary Figure 27. Pore radius of the the heptamer calculated with HOLE2.**[22] Each line corresponds to a 500 ps frame. Right-side shows maximum radius fluctuation over time for the N-terminus (top, pore coordinate approx. 15 Å) and C-terminus (bottom, pore coordinate approx. 35 Å). 3 x 500ns runs from starting poses in Supplementary Figure S24.

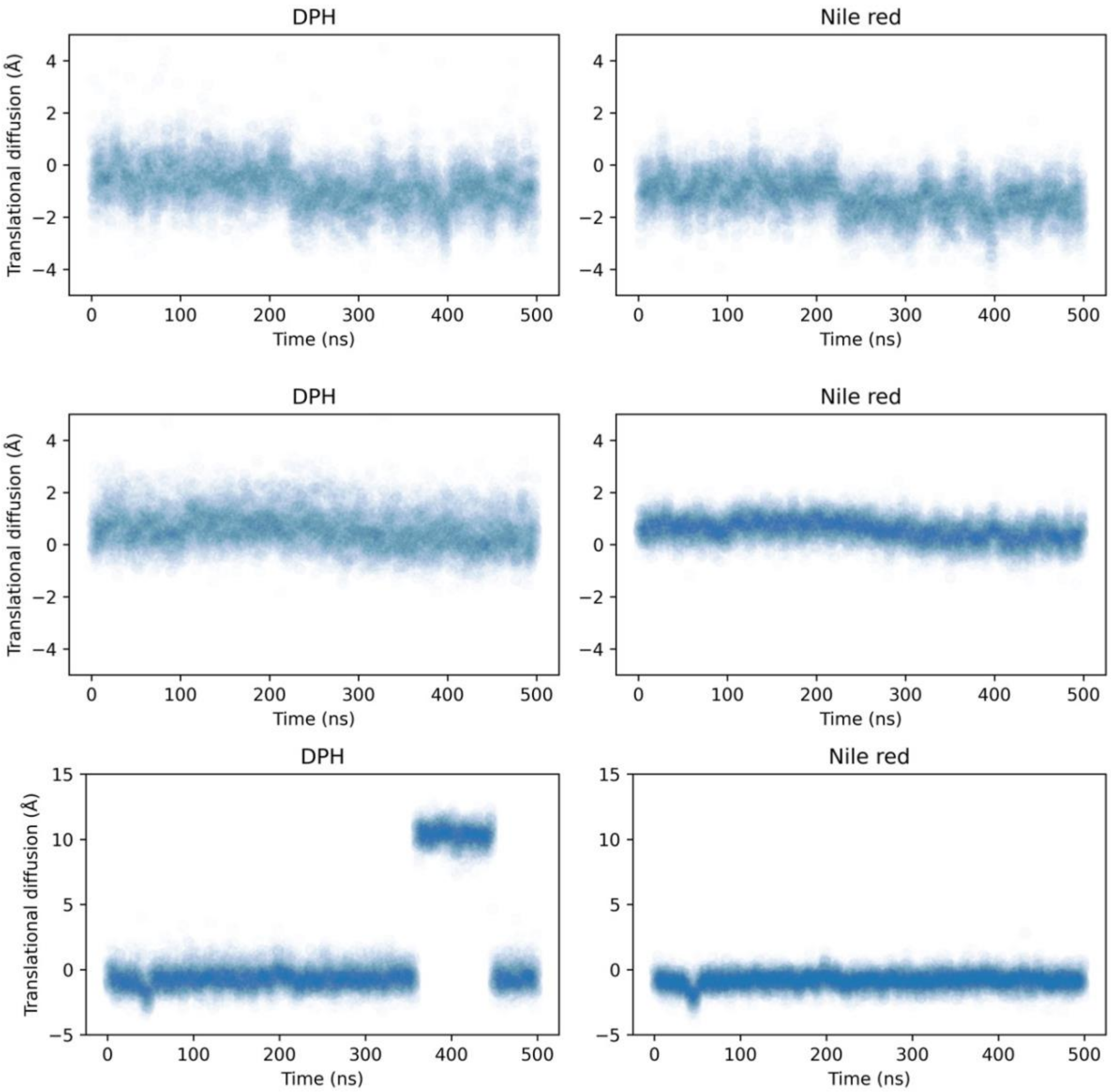

**Supplementary Figure 28. Translational diffusion of DPH and Nile Red within the heptamer along the channel.** 3 x 500ns runs from starting poses in Figure S24.

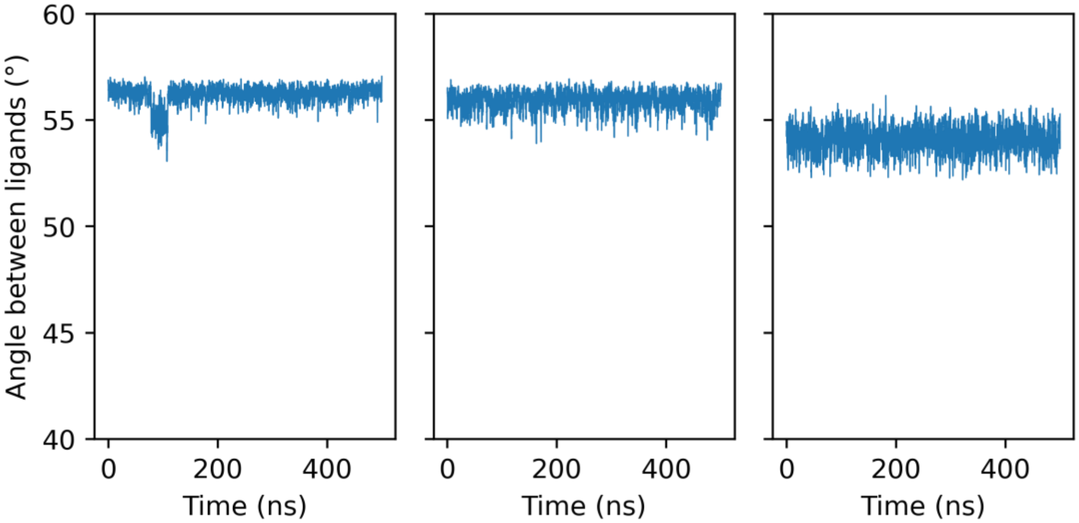

**Supplementary Figure 29. Angle between DPH and Nile red within the heptamer.** 3 x 500ns runs from starting poses in Figure S24. The angle was calculated between the long axes of the molecules.

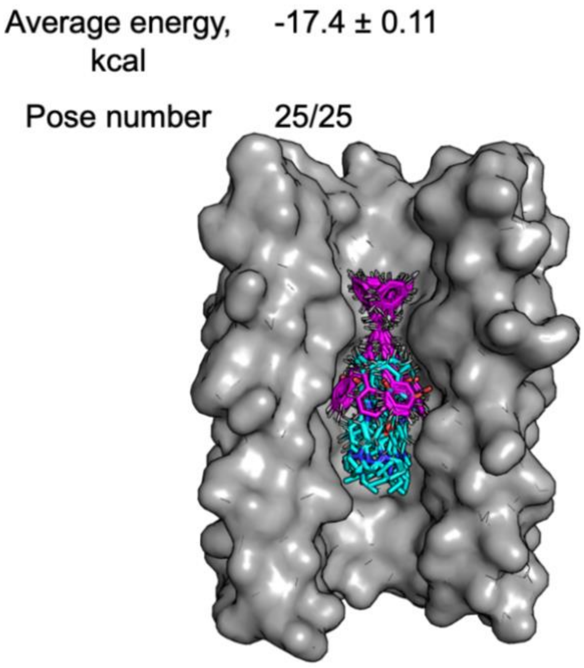

**Supplementary Figure 30. Top 28 AutoDock Vina binding poses for DPH (purple) and Nile Red (cyan) in the octamer (PDB: 9f5a).** Ligands were geometry-optimised with a MOPAC (PM6-D3H4, COSMO) and simultaneously docked with AutoDock Vina 1.2 with exhaustiveness of 64. DPH torsion angles were set to 0.

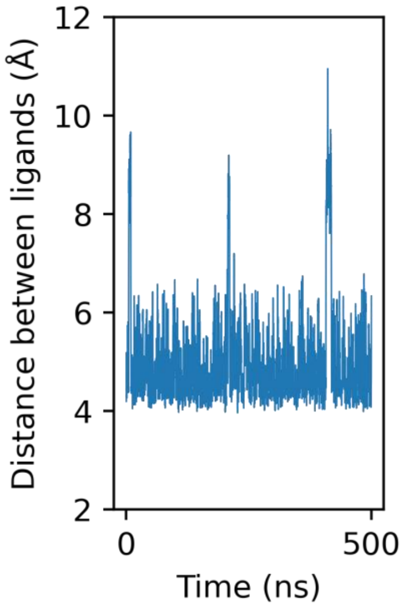

**Supplementary Figure 31. Distance between DPH and Nile red within the octamer.** 500ns simulation from a starting pose in Figure S30. The distance was calculated between the centres of mass of the molecules.

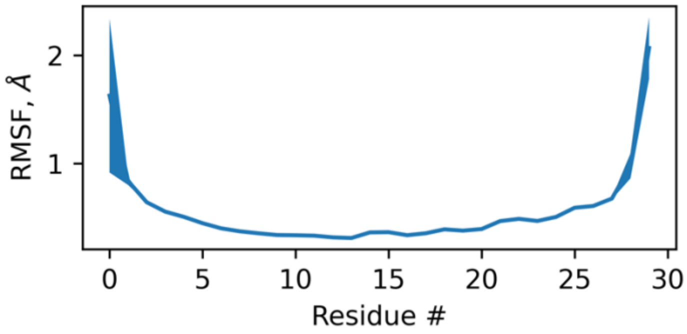

**Supplementary Figure 32. Cα root mean square fluctuation in the octamer.** Line area shows mean across all chains ± one standard deviation. 500 ns simulation from a starting pose in Figure S30.

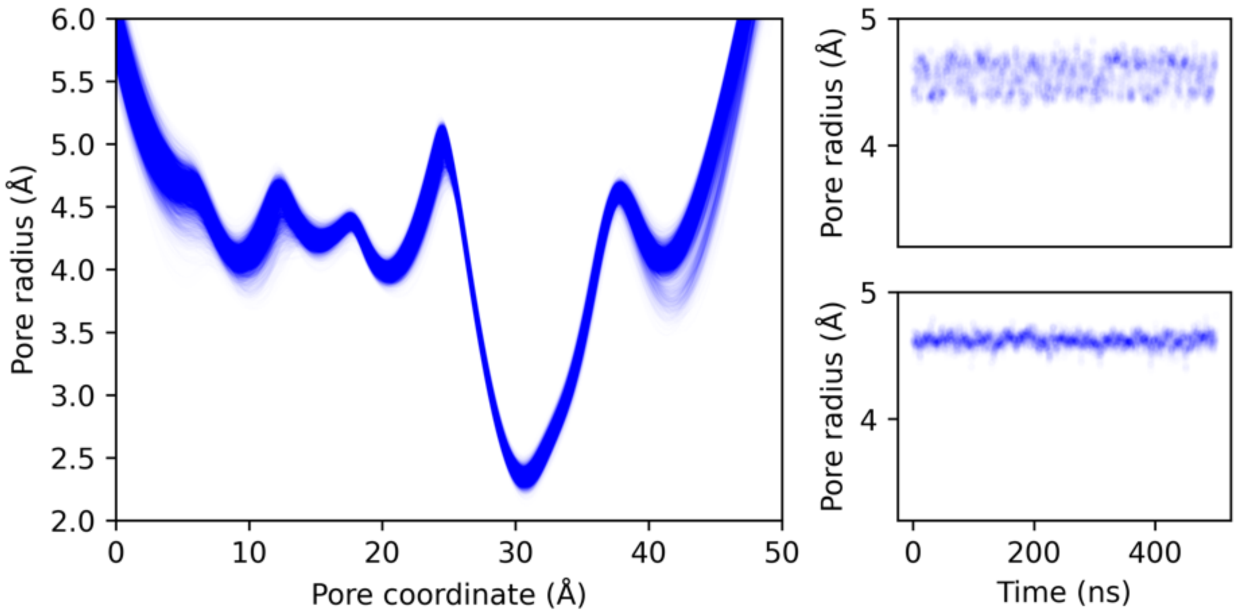

**Supplementary Figure 33. Pore radius of the the octamer calculated with HOLE2.**[22] Each line corresponds to a 500 ps frame. Right-side shows maximum radius fluctuation over time for the N-terminus (top, pore coordinate approx. 15 Å) and C-terminus (bottom, pore coordinate approx. 35 Å). 500 ns simulation from a starting pose in Figure S30.

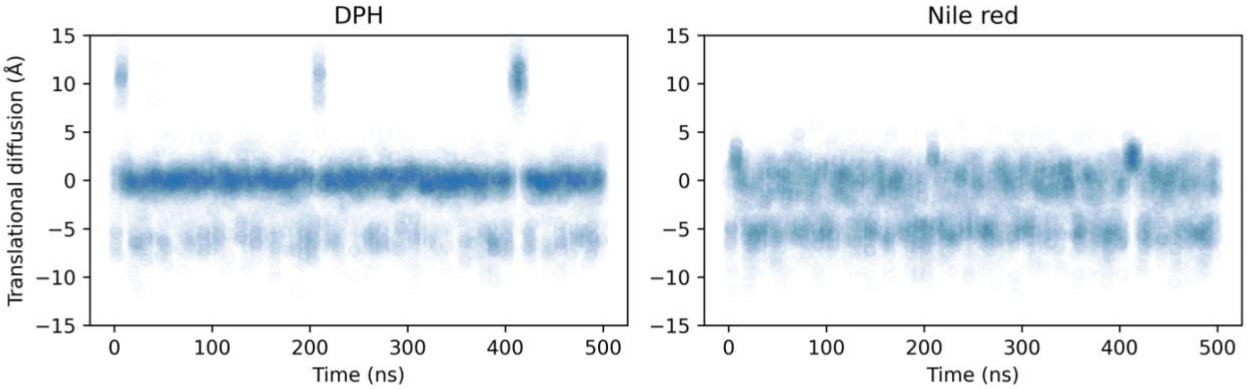

**Supplementary Figure 34. Translational diffusion of DPH and Nile Red within the octamer along the channel.** 500 ns simulation from a starting pose in Figure S30.

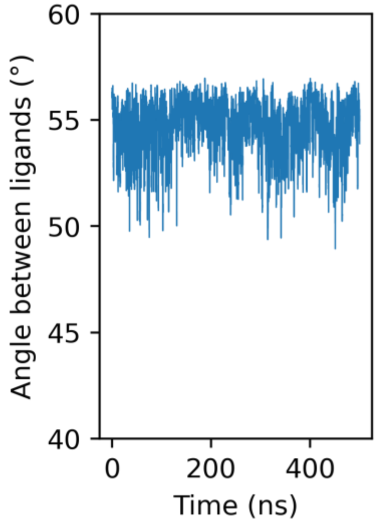

**Supplementary Figure 35. Angle distribution between DPH and Nile Red within the octamer.** 500 ns simulation from a starting pose in Figure S30. The angle was calculated between the long axes of the molecules.

##### 3.7 Anthracene photodimerization

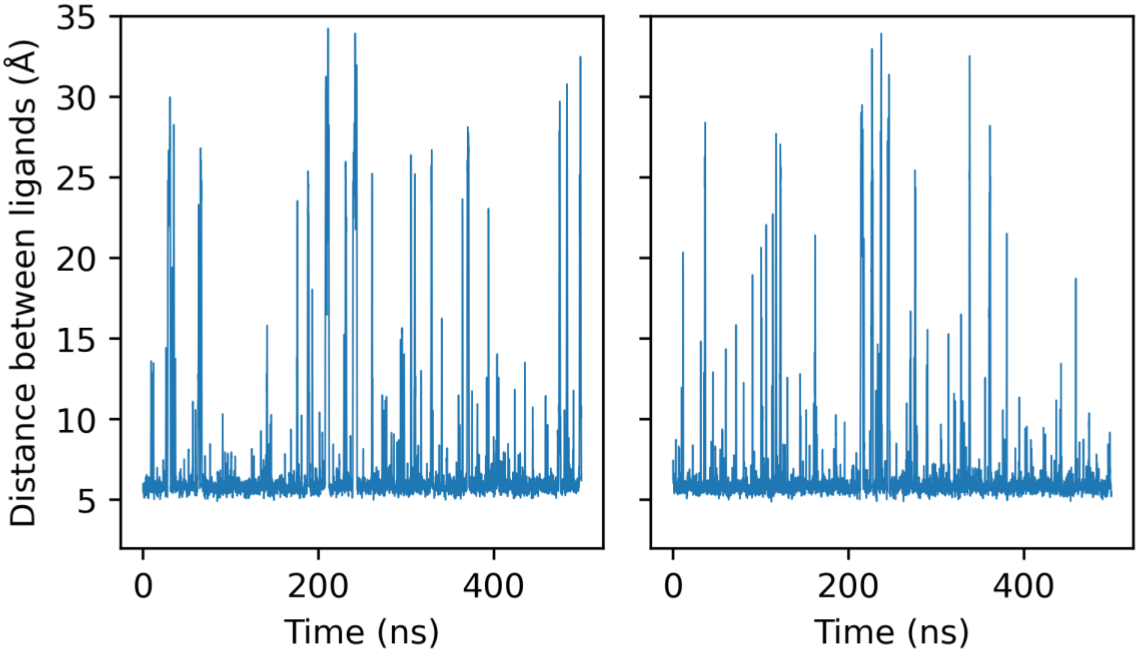

**Supplementary Figure 36. The distance between two anthracene molecules during the 0.5 μs MD simulations initiated from poses in Figure 5a**. Centre of mass was used for calculating the distance.

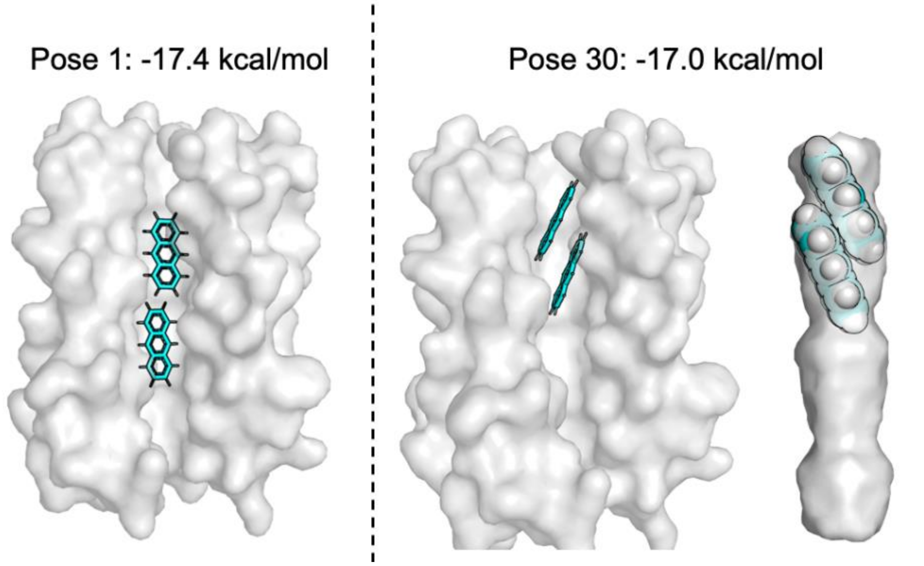

**Supplementary Figure 37. AutoDock Vina binding poses for anthracene in the hexamer.** Top energy pose and a partially stacked pose showing the channel volume. Ligands were geometry-optimised with a MOPAC (PM6-D3H4, COSMO) and simultaneously docked with AutoDock Vina 1.2 with exhaustiveness of 64.

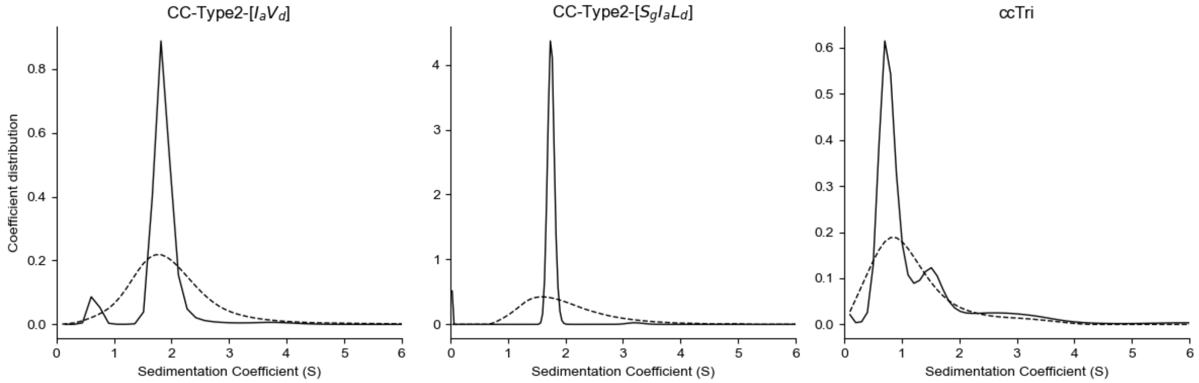

**Supplementary Figure 38. AUC sedimentation velocity data fits (single-species c(s) model with partial peptide volume in solid and ls-g*(s) model in dashed) for anthracene sedimentation within the heptamer, the hexamer and the trimer.** Data collected at 250 nm and 50 000 RPM. Fitted mass: 23.4 kDa for the heptamer, 22.5 kDa for the hexamer**.** Conditions: 15 μM anthracene, 15 μM peptide assembly, HEPES, 10% v/v MeCN, pH 7. Anthracene co-sedimentation with the trimer but lack of the induced CD signal (Figure 5b) is consistent with non-specific binding, similar to that observed with DPH and Nile Red in Figure S7.
